# Diverse modes of T cell receptor sequence convergence define unique functional and cellular phenotypes

**DOI:** 10.1101/2025.05.31.657155

**Authors:** Stefan Schattgen, Kasi Vegesana, William D. Hazelton, Anastasia Minervina, Sebastiaan Valkiers, Kamil Slowikowski, Neal Smith, MGH COVID-19 Team, Alexandra-Chloé Villani, Paul G. Thomas, Philip Bradley

**Affiliations:** Department of Host-Microbe Interactions, St. Jude Children’s Research Hospital, Memphis, TN, USA; Department of Informatics, University of Antwerp, Antwerp, Belgium; Computational Biology Program, Division of Public Health Sciences, Fred Hutchinson Cancer Center, Seattle, WA, USA; Center for Immunology and Inflammatory Diseases, Department of Medicine, Massachusetts General Hospital, Boston, MA, USA; Krantz Family Center for Cancer Research, Massachusetts General Hospital, Boston, MA, USA; Broad Institute of Massachusetts Institute of Technology and Harvard, Cambridge, MA, USA; Harvard Medical School, Boston, MA, USA; Institute for Protein Design, Department of Biochemistry, University of Washington, Seattle, WA, USA

## Abstract

Single-cell techniques allow concurrent study of gene activity and T cell receptor (TCR) sequences, identifying connections between TCR structure and cell traits. Expanding on our CoNGA software, we present a “metaCoNGA” analysis of 6 million T cells from 91 diverse studies, mapping TCR sequence similarity across tissues and diseases. This approach exposes shared TCR features within specific T cell subsets, including those associated with infection, cancer, and autoimmunity. We introduce a method to identify T cell groups with similar gene expression and biased TCR amino acid composition, providing a systematic framework for classifying diverse unconventional T cells, including KIR+ CD8+ T cells, CD4+ regulatory T cells, and subsets of NKT and MAIT cells. A new TCR clustering approach identifies thousands of convergent TCR sequence clusters hypothesized to target shared antigens. These clusters show coherent gene expression, highlighting the role of antigen exposure in shaping T cell behavior. Finally, we provide a tool for users to merge new data with this resource and rapidly identify T cell features in their data sets. This resource empowers investigations into the complex relationship between TCR sequence and T cell function in human health.

## Introduction

The binding properties of the TCR play a fundamental role in determining a T cell’s development and phenotype, from thymic selection to survival and activation in the periphery. These binding properties are determined by the TCR’s amino acid sequence; thus it is not surprising that there are discernable relationships between TCR sequence and transcriptional phenotype. For example, mucosal-associated invariant T (MAIT) cells share TCR sequence features (V and J gene biases, Complementarity Determining Region 3 [CDR3] sequence motifs) that correlate with their distinctive gene expression signatures [1]. TCRs specific for the same pMHC epitope tend to have similar TCR sequences [2,3] and have been observed to have similar phenotypes in many cases [4–8]. Relationships between TCR sequence and cellular phenotype are also present at the level of TCR sequence composition: T cell subsets such as CD4+ T cells or regulatory T cells (Tregs) exhibit TCR sequence composition biases such as elevated positive charge (CD4+ T cells) or enhanced hydrophobicity (Tregs) [9–11]. These TCR sequence compositional biases are modest, and they are superimposed onto the inherent TCR sequence diversity associated with recognition of diverse epitopes, yet they are highly statistically significant when assessed at the level of large T cell populations.

Whereas the discovery of these relationships between TCR sequence and cellular phenotype required prior knowledge of the relevant T cell populations, the recent application of multimodal single-cell technologies to large populations of T cells derived from blood and tissue of diverse donors has produced data resources [12–15] that enable unbiased and systematic exploration of TCR-GEX correlation. With such data and sophisticated algorithms, it may even become possible to model the mapping between TCR sequence and cellular phenotype, at the population level or conditioned upon an individual’s HLA genotype and immune exposures. As a step toward this goal, we assembled published and unpublished multimodal T cell datasets encompassing 6 million T cells with paired TCR sequences (4 million unique TCR clonotypes), from 91 studies on infection, cancer, autoimmunity, and aging across a range of tissues (the ‘metaCoNGA atlas’, **Fig. 1** and **Table S1**).

**Figure 1.**
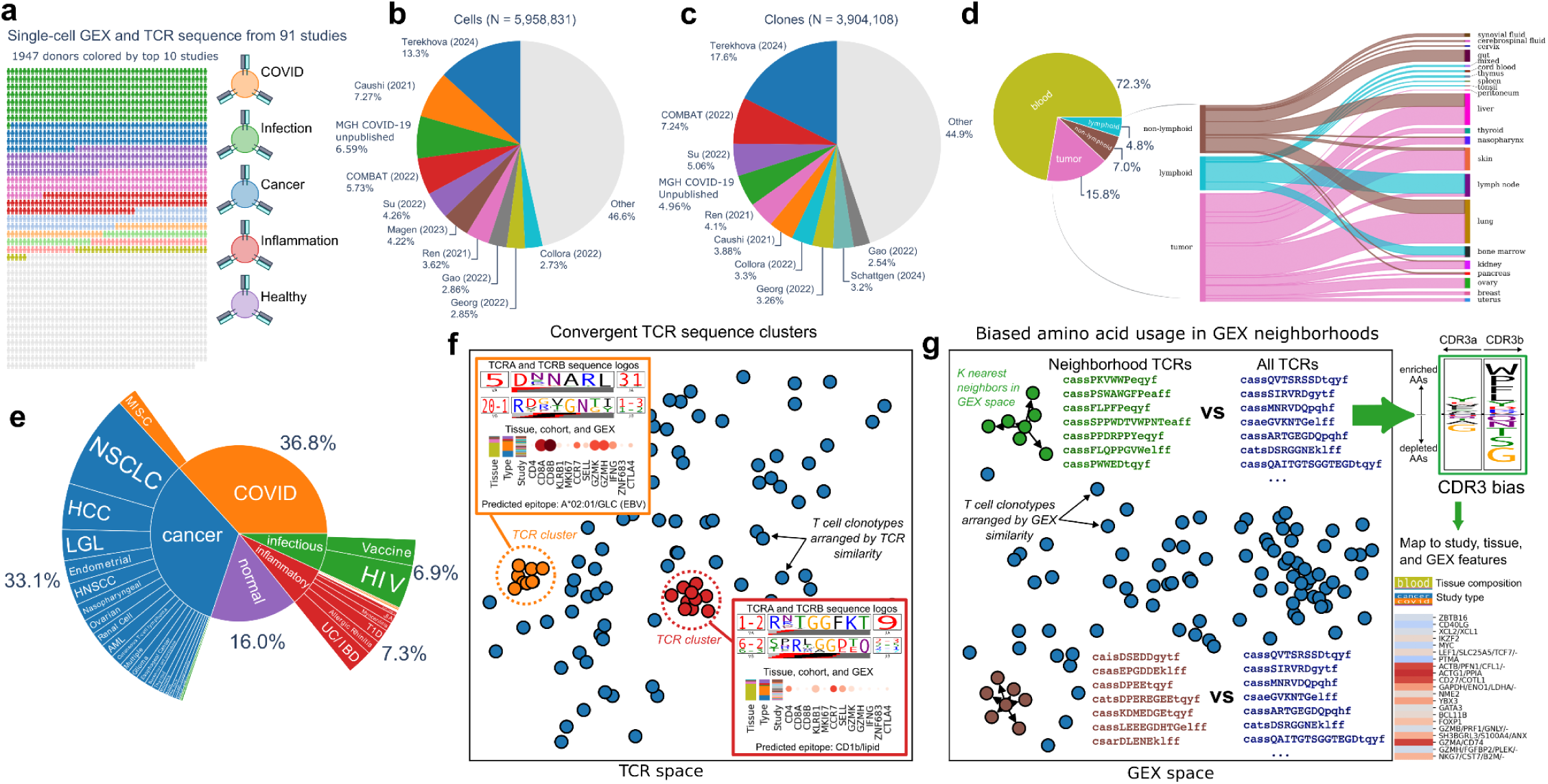
The metaCoNGA dataset. (a) Ninety-one single-cell GEX and TCR repertoire datasets generated using the 10X Immune Profiling assay were identified, retrieved, processed, and integrated with only T cells with paired TCRαβ information included. (b) Pie chart showing the ten largest studies in the dataset by number of cells. (c) Pie chart showing the ten largest studies in the dataset by number of unique paired TCR sequences. (d) Pie chart (left) and a Sankey plot showing the specific tissues of origin by cell frequency. (e) Sunburst plot of the disease conditions at broad and refined levels associated with samples included in the dataset. (f) Schematic landscape of global TCR sequence similarity among T cell clonotypes (filled circles), with two convergent clusters highlighted in red and orange and mapped to their TCR sequence features (logos) and gene expression, tissue, and source study correlates. (g) Landscape of GEX similarity among T cell clonotypes showing two KNN neighborhoods with biased CDR3 sequence composition and phenotypic features.

With this multimodal collection assembled, we looked systematically for human T cell populations exhibiting global and compositional sequence convergence. We extended the graph-based algorithms in the CoNGA software package [4] to perform a statistical analysis of TCR similarity and its relationship to gene expression in this merged dataset, identifying multiple modes of TCR sequence convergence that correlate with cellular phenotype (**Fig. 1f-g**). First, we developed a TCRdist-based [2] clustering algorithm to identify and group convergent TCRs using a robust, data-derived background model **(Fig. 1f)**. This method identified more than 100,000 TCR clonotypes with statistically significant global sequence convergence; these clonotypes can be grouped into clusters of which ∼2,200 contain at least 10 clonotypes **(Fig. 2)**. By matching these TCR sequences to databases of known antigen-specific TCRs, as well as TCRs associated with CMV serostatus and specific HLA alleles, we were able to assign putative epitope and HLA restrictions to many of the TCR clusters. Notably, the GEX profiles within these sequence-based clusters were highly coherent, and TCR clusters specific to antigens from the same pathogen showed greater similarity to one another than to those from unrelated pathogens. The second approach we developed aims to detect graph neighborhoods in GEX space that exhibit compositional sequence convergence by examining for biased amino acid usage within the hypervariable CDR3 **(Fig. 1g)**. This approach found that 9% of CD8+ and 2% of CD4+ GEX neighborhoods showed statistically significant amino acid sequence bias at a stringent false discovery threshold **(Figs. 3-5)**. Some of these compositionally-biased cell populations map to known unconventional T cell populations such as KIR+ CD8+ T cells, CD4+ Treg cells, and diverse subsets of NKT and mucosal-associated invariant T (MAIT) cells. Additional subpopulations were found to be specifically enriched in contexts of cancer and autoimmunity. A key finding of our analysis is that both global and compositional sequence convergence are widespread and are deterministic in shaping cellular phenotypes. Taken together, these findings suggest that multimodal single-cell genomics data on T cells, coupled with advanced algorithms, will significantly advance our understanding of human immunology.

**Figure 2.**
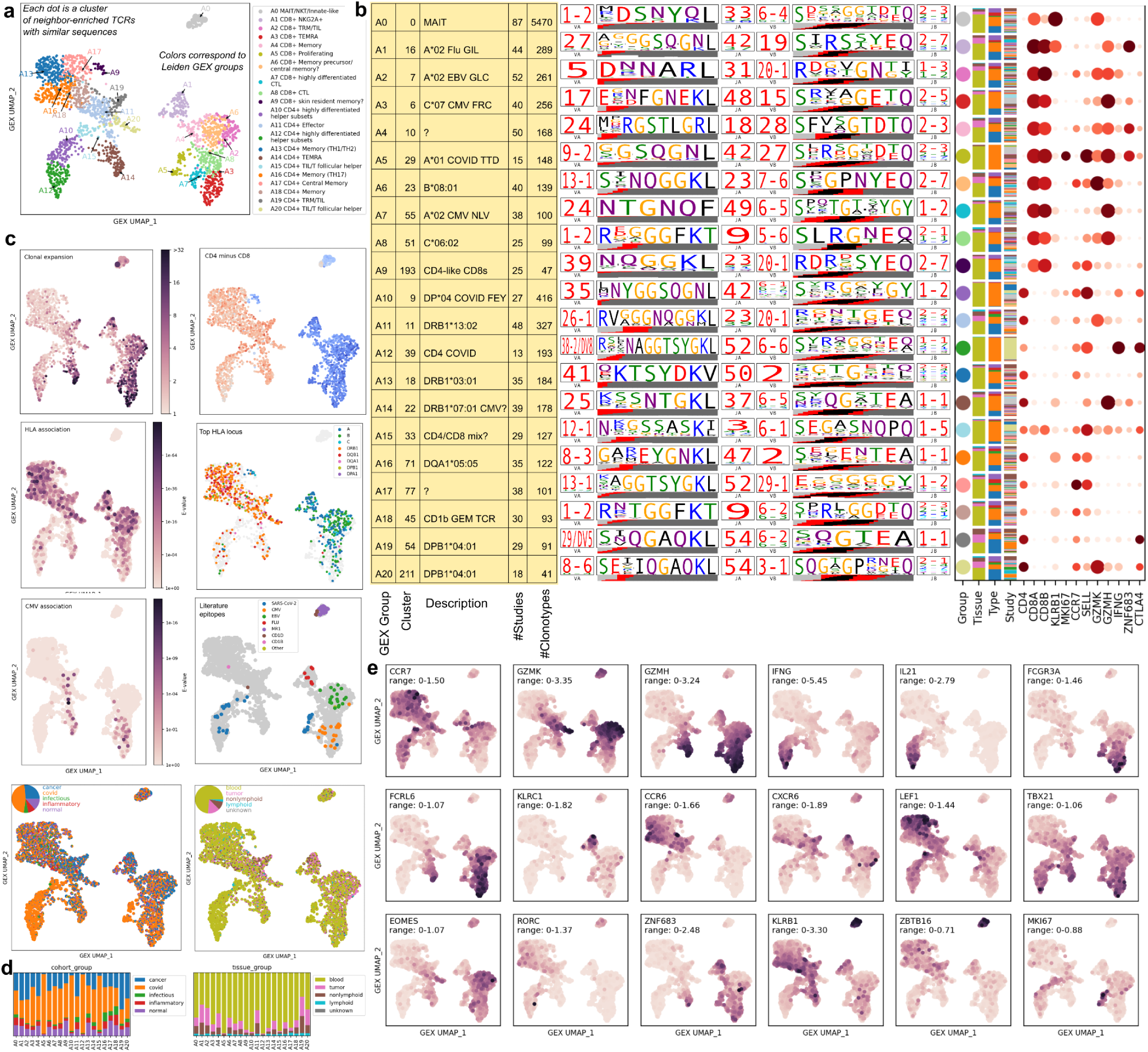
Landscape of TCR sequence convergence. (a) 2D GEX landscape of convergent TCR sequence clusters. Colors represent 21 GEX-derived groupings of the individual clusters; the largest cluster in each group is indicated with an arrow and shown in the sequence logos to the right. (b) Annotations, sequence logos, and marker gene expression for the largest TCR cluster in each GEX group. (c) Panels showing the 2D GEX landscape colored by clonal expansion, CD4-CD8 expression difference, HLA- and CMV-associations, sequence matches to TCR literature databases, study type and tissue. (d) Study type and tissue breakdown for the 21 GEX groupings of the clusters. (e) Marker gene expression averaged over the cells of each TCR cluster and mapped onto the 2D GEX landscape.

**Figure 3.**
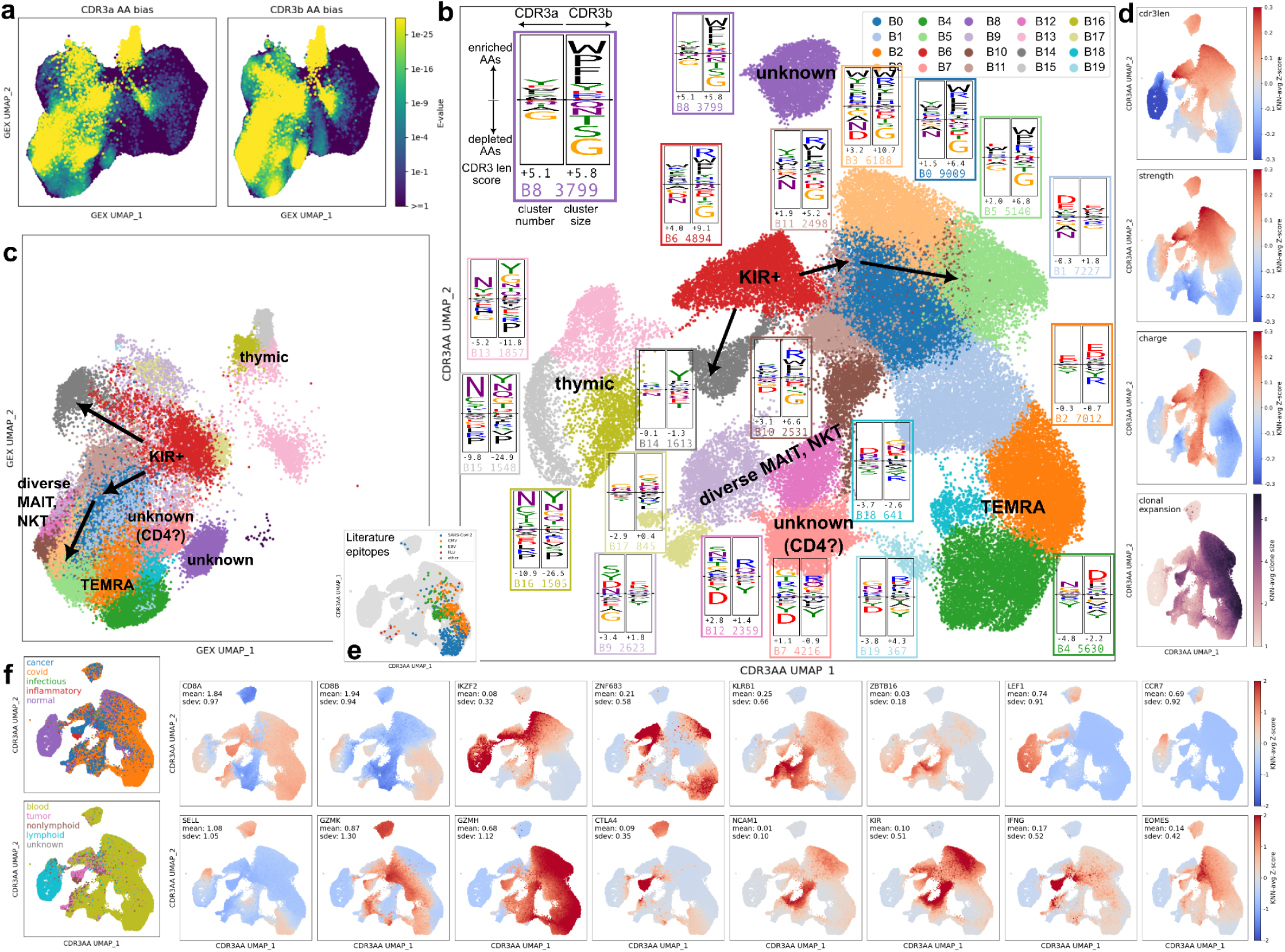
CDR3 amino acid biases in CD8+ GEX neighborhoods. (a) 2D GEX landscape colored by the multiple-testing-corrected significance score for each neighborhood (represented by the central clonotype; CDR3α scores on the left and CDR3β scores on the right). (b) Each significantly biased neighborhood was mapped to its forty-dimensional vector of observed amino acid (AA) frequencies (20 for CDR3α and 20 for CDR3β); these frequency vectors were Leiden clustered (colors) and mapped into 2 dimensions via the UMAP algorithm. Next to each cluster is a differential sequence logo showing enriched and depleted amino acids, CDR3α and CDR3β length scores (positive for longer than average CDR3s, negative for shorter than average), and the cluster number and size. (c) The same CDR3AA-biased neighborhoods mapped to the GEX landscape to show the consistency of the frequency-vector-derived clusters (colors). (d) CDR3 length, strength, charge, and clonal expansion mapped onto the CDR3AA frequency landscape. (e) Statistically significant sequence matches to TCRs from the literature. (f) Study type, tissue type, and marker gene expression mapped onto the CDR3 AA frequency landscape.

## Results

### MetaCoNGA Dataset

To explore relationships between T cell receptor (TCR) sequence and cellular phenotype, we combined 91 human single-cell datasets from >1900 subjects that featured paired TCR alpha/beta sequence and mRNA transcript counts (**Fig. 1a**). The merged dataset contains ∼6 million cells and ∼4 million unique TCR clonotypes (**Table S1**, **Fig. 1b+c**) distributed across several larger cohort studies and many smaller disease- and organ-focused datasets. By consulting the original studies and associated metadata, we were able to assign a tissue of origin to >99% of the cells. Grouping these tissue assignments into broad categories yields the breakdown shown in **Figure 1d**: 72.3% peripheral blood mononuclear cells (PBMC), 15.8% tumor, 7.0% non-lymphoid (kidney, liver, lung, etc), 4.8% lymphoid (bone marrow, spleen, thymus, tonsil, lymph node, and cord blood), and the remainder were samples mixed from multiple tissues. Classifying the individual studies into five broad groups shows that the two largest groups, COVID19 and cancer, make up 36.8% and 33.1% of the dataset, respectively (**Fig. 1e**). Samples from healthy individuals (16%), auto-inflammatory/immune diseases (7.3%), and non-COVID19 infection or vaccination (6.9%) settings account for the remainder (**Fig. 1e**). To facilitate merging these diverse datatsets, we developed a new approach to feature selection that leverages the paired TCR sequence data. Rather than the standard approach of selecting highly variable genes (HVGs) as features for downstream analysis, we identified genes that varied significantly across the TCR neighbor graph (**Table S2**), using a graph autocorrelation statistic [16] (see Methods). We observed that these TCR-graph autocorrelated genes (TAGs) encompassed many of the key T cell marker genes and generated more consistent and interpretable landscapes and clusterings than HVG-based feature sets (**Figs. S1-2**). Using this dataset, we identified correlations between TCR sequence and gene expression across a spectrum ranging from global TCR sequence similarities (**Fig. 1f**)—likely driven by shared antigen recognition—to CDR3 sequence composition biases (**Fig. 1g**) characteristic of specific T cell subpopulations. Our objective was to uncover these correlations in an unbiased manner, enabling the discovery of unexpected cell populations and novel TCR-GEX relationships.

### TCR sequence convergence

To assess global TCR sequence convergence in the dataset, we used the TCRdist distance metric [2] and a realistic background model of V(D)J-rearrangement (see Methods) to identify TCR clonotypes with a greater than expected number of sequence neighbors (see Methods, **Fig. S3**). Inspired by the ALICE algorithm [17], we assign a Poisson *p*-value to each clonotype that reflects our surprise at seeing the observed number of TCR sequence neighbors in the dataset given an expected neighbor rate for that clonotype estimated from the background. Applying a stringent Bonferroni correction that accounts for the total number of clonotypes, we identified 109,384 neighbor-enriched TCR sequences (∼2.8% of the dataset). We grouped these neighbor-enriched TCRs into clusters of similar sequences (see Methods), of which 2,173 had at least 10 members. Using the top 200 TAGs (**Fig. S1b**) as features, gene expression was averaged over all members of each cluster prior to performing PCA, Leiden clustering, and UMAP dimensionality reduction to create a gene expression landscape of convergent TCR clusters (**Fig. 2a**; gene expression was strikingly coherent within clusters, **Figs. S4-5**). Manual inspection of marker genes for each of the 21 Leiden GEX groupings (A0-A20) demonstrated a spectrum of identifiable differentiated (i.e. non-naive) T cell subtypes (**Fig. 2a**; **Table S3**). Antigen-driven selection is a dominant driver of TCR sequence convergence and explains the notable lack of naive T cells detected in this analysis. TCR sequence logos and marker genes for the largest cluster in each GEX grouping are shown in **Figure 2b**. Coloring the landscape by the CD4/CD8 gene expression difference (**Fig. 2c**) shows a population of CD8+ clusters on the right and CD4+ clusters on the left (with a small CD8 “island” nearby). The gray clusters at the top of **Figure 2a**, marked by expression of *KLRB1* and *ZBTB16* in **Figure 2e**, contain MAIT and iNKT TCRs (TRAV1-2/TRAJ33 gene usage in the top sequence logo in **Figure 2b**). In the CD8 clusters, a GZMK/GZMH gradient is apparent, with GZMK higher in clusters near the top and GZMH higher in clusters near the bottom. Cytotoxic CD4 T cells can be seen on the right-hand side of the CD4 region, closest to the CD8 clusters, and the same GZMK/GZMH gradient is visible in these clusters.

Matching these clusters to TCR sequences from the literature (see Methods) highlights a tendency for T cells responding to the same pathogen to have similar gene expression, with CMV-, EBV-, FLU- and SARS-CoV-2-reactive T cells generally being found in distinct regions of GEX space (**Fig. 2c**). In particular, CMV-reactive CD8+ T cells are found in the *GZMH*-hi region of the landscape whereas EBV-reactive T cells tend to be found in the *GZMK*-hi region. We assigned putative HLA restriction and CMV-association to many of these convergent TCR clusters (**Fig. 2c**) by tracking occurrences of their TCR beta chains in a large set of bulk repertoires from HLA-typed individuals of known CMV serostatus [18]. The CMV associations support the CD8+ CMV landscape boundaries derived from literature matching and define additional putative CD4+ CMV-reactive sequences (e.g., in GEX group A14, colored brown in **Fig. 2a**). The majority of the HLA assignments are consistent with the CD4/CD8 status of the clusters (**Fig. 2c**), with HLA class I assignments seen on the right-hand side of the landscape in the clusters of CD8+ T cells and HLA class II assignments observed within the CD4+ clusters. Looking into examples of clusters with inconsistent top HLA associations (class II associations for CD8+ clusters or class I for CD4+ clusters), many have a slightly weaker but still significant association with an HLA class-matched allele (for example, highly significant associations with both A*01:01/B*08:01/C*07:01 and DRB1*03:01/DQA1*05:01/DQB1*02:01, due to their tendency to co-occur as a haplotype). A number of GEX groups showed a skewed composition in terms of tissue and disease relative to their global frequencies. GEX groups A11, A19, and A20, for example, were enriched with T cells from tumor samples which displayed phenotypes consistent with CD4+ effector, tissue resident memory, and follicular helper cells, respectively, that likely correspond to responses associated with tertiary lymphoid structures in tumors [19].

### Biases in CDR3 sequence features

Identification of TCR sequence biases characteristic of specific T cell subpopulations, such as CD4+ T cells or Tregs, has traditionally required advanced knowledge of the relevant populations to enable their isolation and sequencing. Additionally, it has relied on advance knowledge of the biased amino acid features (e.g., CDR3 length, charge, hydrophobicity) to assess their distributions. To take an unbiased look at CDR3 sequence biases in phenotypically-defined subpopulations, we looked for GEX graph neighborhoods (i.e., a central clonotype together with its K nearest neighbors in gene expression) whose CDR3 amino acid (CDR3AA) frequency distributions deviated significantly from the background amino acid distribution for the entire dataset (see Methods and **Fig. S6**). Given that CD4+ and CD8+ T cells are known to have different CDR3AA frequencies, we performed this analysis over the CD4+ and CD8+ subsets separately. We also excluded the large MAIT cluster (upper left cluster in **Fig. S1c**) from the CD8+ dataset, to prevent the strong sequence signal of invariant TCR chains from skewing the background distribution and swamping other features. We assessed the significance of the deviation between each neighborhood’s observed amino acid frequencies and the expected background frequencies using the chi-squared distribution, corrected for the total number of neighborhoods analyzed. The size of our dataset allowed for the use of large neighborhoods (K=5000 nearest neighbors, 0.2% and 0.6% of the CD4 and CD8 datasets, respectively) to maximize statistical power without sacrificing neighborhood locality.

We found 71,528 CD8+ GEX neighborhoods (8.7% of all CD8+ neighborhoods) that showed highly significant CDR3AA sequence biases (Bonferroni-corrected *p*-values less than 10^−6^; **Fig. 3a**). We projected each significant neighborhood to a 40-dimensional amino acid frequency vector (20 CDR3α frequencies and 20 CDR3β frequencies) and performed UMAP dimensionality reduction and Leiden clustering on these frequency vectors to visualize this landscape of CDR3AA bias (**Fig. 3b**). Each of the 20 Leiden CDR3AA clusters (B0-B19) in **Figure 3b** is overlaid with the amino acid frequency logo for its CDR3α (left) and CDR3β (right) loops, with overrepresented amino acids shown above the logo midpoint and depleted amino acids shown below the midpoint. Although the CDR3AA clusters are defined entirely based on CDR3 amino acid frequencies, they remain largely coherent when their central clonotypes are projected back into GEX space (**Fig. 3c**). Plotting selected TCR features averaged over these significantly biased neighborhoods showed heterogeneity in CDR3 length, strength, and charge across the embedding (**Fig. 3d**). In the upper half of the CD8+ CDR3AA bias landscape are seven clusters (B0, B3, B5, B6, B8, B11 and B14) with high interaction strength (approximated by enrichment for specific ‘strongly-interacting’ amino acids; **Fig. S1g**), generally longer CDR3 loops, and higher charge (**Fig. 3b+d**). Their sequence logos show overrepresentation of large amino acids such as tryptophan, tyrosine, and phenylalanine. Clusters B1, B2, B4, and B18 on the lower right of the landscape have lower than average CDR3 length, interaction strength, and charge (**Fig. 3b+d**). They show higher expression of cytolytic genes including granzymes and granulysin, and they have a higher fraction of neighborhood clonotypes belonging to convergent TCR clusters (**Fig. S7a**, ‘is_convergent’ panel) including CMV, EBV, and SARS-CoV2-specific TCRs (**Fig. 3e+f**). Clusters B13, B15, and B16 (left side of the landscape) contain thymic T cells whose distinctive sequence features (e.g. enrichment for tyrosine and asparagine, shorter CDR3 lengths) and marker genes (e.g. *LEF1* and *CCR7*) likely reflect their unique developmental stage (**Fig. 3b+d+f**; top differentially expressed genes for all clusters shown in **Fig. S8**).

We performed targeted GEX analysis (see Methods) on the CD8+ neighborhoods with biased CDR3AA composition, yielding a focused GEX landscape and 17 GEX clusters (C0-C16) whose structure was largely coherent with the CDR3AA-derived landscape (**Fig. 4a-c**). GEX clusters C10 and C13 were composed of the thymic T cells from CDR3AA clusters B13, B15, and B16, and expressed the transcription factors *SOX4* and *IKZF2*, and chemokine receptor *CCR9*, consistent with agonistically-selected thymic T cell subsets [20–23] (**Fig. 4d**). GEX clusters C1-3, C6, C8, and C14 (CDR3AA clusters B0, B1, B3, B5, B6, and B11) exhibited GEX features similar to the recently-described *KIR+* CD8 T cells [4,24–28]. Substantial diversity is evident, however: CDR3AA cluster B6 is most similar to the *KIR+* population identified in our earlier work [4], being positive for both *ZNF683* and *IKZF2* genes, only modestly clonally expanded, positively charged in CDR3 regions, and found primarily in blood. CDR3AA clusters B0, B3, and B5 show increasing clonal expansion (**Fig. 3d**), expression of cytolytic genes (i.e. *GZMB*, *GZMK*), NK receptors (*KLRC3*, *NCR3*) and loss of either *ZNF683* (B0 and B3) or *IKZF2* (B5) (**Fig. 4d**). CDR3AA clusters B1, B2, B4, and B18, marked by antigen-specific responses to CMV, EBV, and SARS-CoV-2 (**Fig. 3e**), occupied GEX clusters C0, C7, and C12 and expressed *GZMB*, *GZMK, ZNF683,* and *FCGR3A*, consistent with terminally differentiated effector memory T cells (TEMRA). GEX clusters C4, C5, and C11 (CDR3AA clusters B7, B9, B10, B12, B17, and B20) expressed markers consistent with non-classical MHC-restricted diverse NKT (dNKT) and MAIT (dMAIT), and killer innate-like T cells (ILTCks) populations including *ZBTB16*, *KLRB1*, *TYROBP*, *FCER1G*, and *CD40LG* [29,30]. GEX cluster C15 (CDR3AA cluster B14) was enriched for tumor infiltrating lymphocytes (TILs) expressing *CXCL13*, *IFNG*, *ENTPD1*, and *GZMB*. CDR3AA cluster B8 was enriched for blood T cells from cancer patients and mapped to GEX cluster C8, which was positive for *GZMK*, *IL10*, and *CXCR5* expression. GEX cluster C16 contained T cells from multisystem inflammatory syndrome in children (MIS-C) patients (CDR3AA cluster B19), including an enrichment in *TRBV11-2* usage, and was positive for *KLRB1* and *GZMB*.

**Figure 4.**
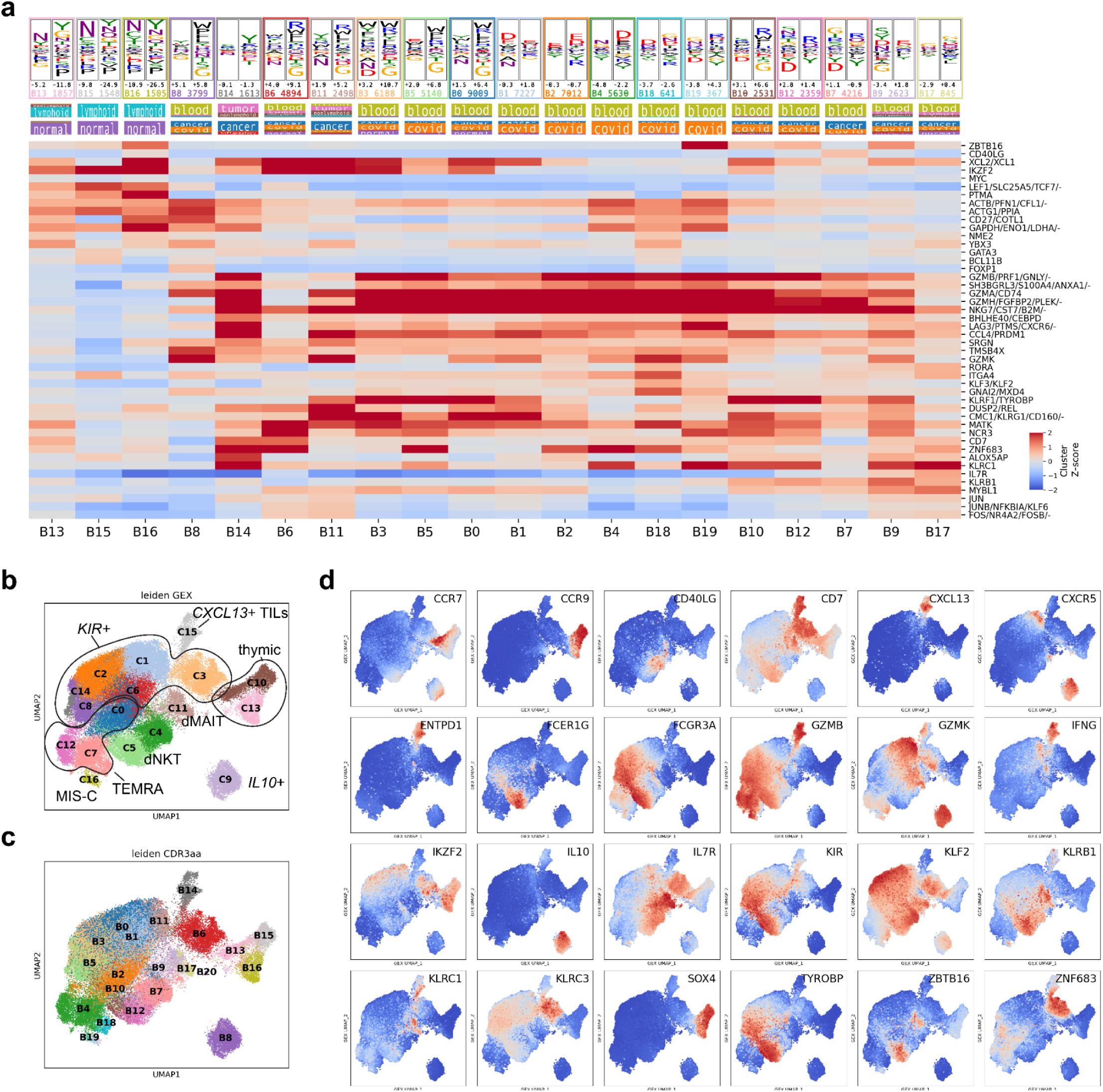
Gene expression analysis of CD8+ neighborhoods with biased CDR3AA composition. (a) Heat map of differentially-expressed genes for the 20 CDR3AA clusters (B0-B19). Highly correlated genes (Pearson R>0.8) were grouped as shown in the labels on the right-hand side. Columns are arranged based on hierarchical clustering using expression levels of genes shown. (b) 2D GEX landscape of CDR3AA-biased CD8+ neighborhoods colored by GEX clusters (C0-C16). (c) 2D GEX landscape of CDR3AA-biased CD8+ neighborhoods colored by CDR3AA clusters (B0-B19). (d) 2D GEX landscape of CDR3AA-biased CD8+ neighborhoods showing indicated GEX features of interest.

Applying a similar analysis to the CD4+ subset of the atlas yielded 53,255 significantly-biased CD4+ neighborhoods (**Fig. 5a**, 2.0% of all CD4+ neighborhoods) that formed 17 CDR3AA clusters (D0-D16) with distinct amino acid composition, biophysical properties, and tissue and disease associations (**Fig. 5b-e**). Based on expression of *FOXP3*, *IKZF2*, and other marker genes (**Fig. 5f**; **Fig. S9**), clusters D0-4, D6, D10, D12, and D13 represent CD4+ Treg cells, with some clusters (D0, D1, D3, D10, D12) found predominantly in blood while others (D2, D4, D6, and D13) are present in tumors and other tissues (**Fig. 5e+f**). The CDR3 regions of the Treg clusters were enriched for hydrophobic residues and depleted for polar or charged residues. One cluster of thymic T cells (D5) expressing *IKZF2* and *CCR7* shared a similar enrichment of tyrosine and asparagine as the thymic clusters in the CD8+ analysis. Four clusters were enriched for MAIT TCR chains (D7, D8, D11, and D14) and expressed markers consistent with cytotoxic innate-like lymphocytes (i.e. *KLRB1*, *ZBTB16*, *CCL5*, *TYROBP*) (**Fig. 5e+f**, **Fig. S7b**, **Fig. S9**).

**Figure 5.**
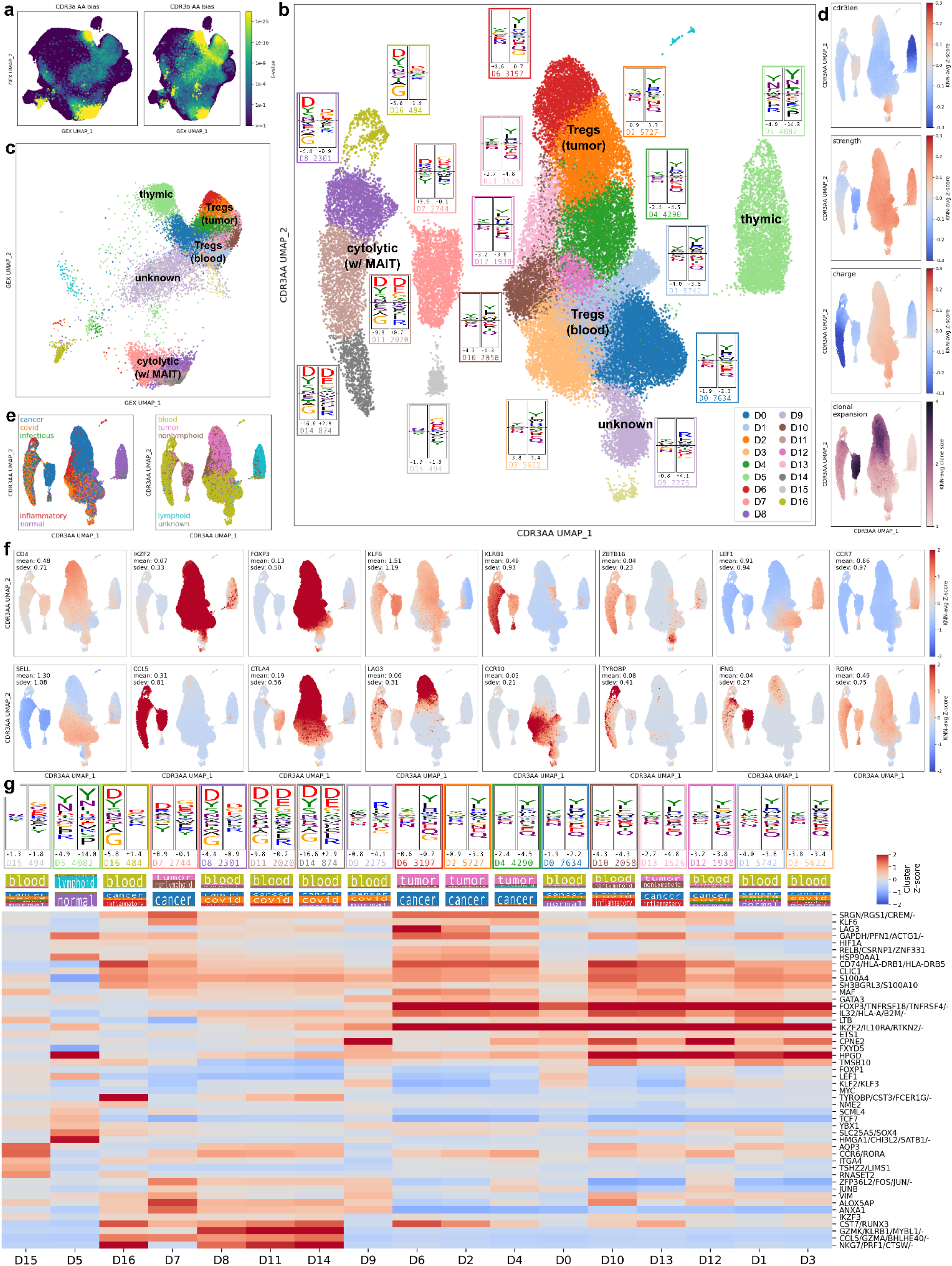
CDR3 amino acid biases in CD4+ GEX neighborhoods. (a) 2D GEX landscape colored by the multiple-testing-corrected significance score for each neighborhood (represented by the central clonotype; CDR3α scores on the left and CDR3β scores on the right). (b) Each significantly-biased (Bonferroni-corrected *p-*values less than 10^−6^) neighborhood was mapped to its forty-dimensional vector of observed amino acid (AA) frequencies (20 for CDR3α and 20 for CDR3β); these frequency vectors were Leiden clustered (colors) and mapped into 2 dimensions via the UMAP algorithm. Next to each cluster is a differential sequence logo showing enriched and depleted amino acids, CDR3α and CDR3β length scores (positive for longer than average CDRs, negative for shorter than average), and the cluster number and size. (c) The same CDR3AA-biased neighborhoods mapped to the GEX landscape to show the consistency of the frequency-vector-derived clusters (colors). (d) CDR3 length, strength, charge, and clonal expansion mapped onto the CDR3 AA frequency landscape. (e) Study type and tissue type mapped onto the CDR3AA frequency landscape. (f) Marker gene expression mapped onto the CDR3AA frequency landscape. (g) Heat map of differentially-expressed genes for the 17 CDR3AA clusters. Highly correlated genes (Pearson R>0.8) were grouped as shown in the labels on the right-hand side. Columns are arranged based on hierarchical clustering using expression levels of genes shown.

### Annotating new datasets with metaCoNGA features

We hypothesized that the convergent TCR clusters and CDR3AA bias motifs identified in the metaCoNGA atlas could help annotate new datasets at the smaller scale of individual donors or samples. To match new TCR sequences to convergent TCR clusters, the TCRdist metric and a background model of V(D)J recombination is used to assign a *p-*value to observed sequence similarity between a TCR in the dataset and a TCR in one of the convergent TCR clusters, allowing us to identify statistically significant matches and thereby transfer annotations such as epitope specificity, MHC restriction, CMV association, etc. This concept is illustrated for individual donor datasets from a recent study of COVID19 infection [31] in **Figure 6**. TCR sequences from one donor that match TCR clusters in GEX group A3 (colored red in **Figs. 2a** and **6b**) are shown in panel 6a along with annotations transferred from the matched clusters, from which we can infer that the donor is likely CMV-positive and positive for the C*07:02 HLA allele. TCR sequences from another donor matching to GEX group A5 are shown in panel 6b; based on source studies and literature matches for A5 clusters, these TCRs are likely COVID19-reactive, with specific HLA and epitope restriction inference possible for a subset. Bar plots represent the fraction of TCR sequences that match TCR clusters, colored by the GEX group (A0-A20) of the matched cluster (**Fig. 6c**) or restricted to matches to COVID19-associated clusters (see Methods) and colored by MHC class restriction (**Fig. 6d**). Notably, the patients in the ‘severe’ disease group (left three bars in **Fig. 6d**) had a higher frequency of matches to COVID19-associated clusters than those in the ‘asymptomatic’ or ‘control’ groups (middle and right bars).

**Figure 6.**
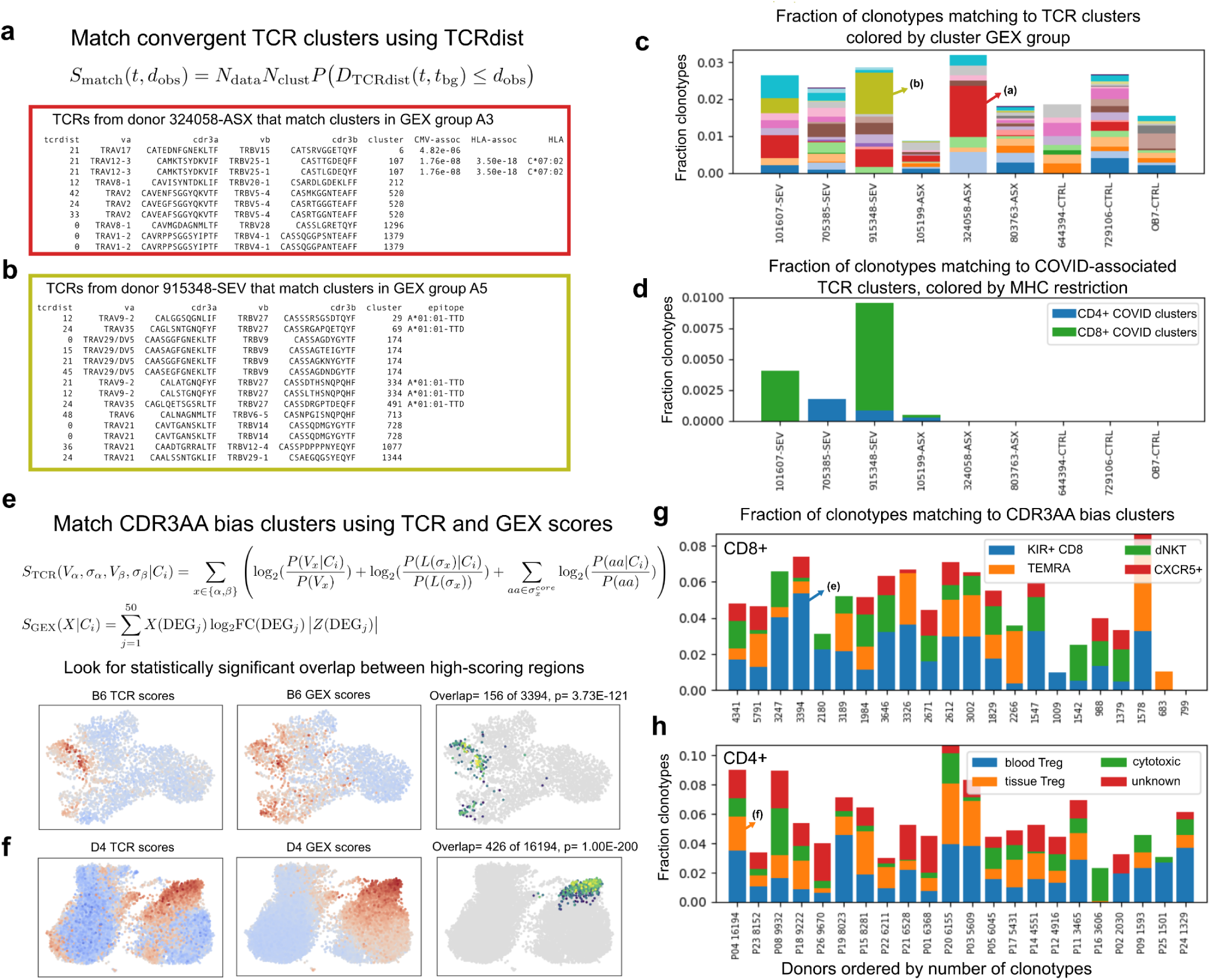
Annotating new datasets with metaCoNGA features. (a) TCRs (*N=N*_data_) in a new dataset are compared to TCRs in the metaCoNGA convergent clusters (*N=N*_clust_) using the TCRdist metric. An observed TCRdist distance (*d*_obs_) between a TCR in the dataset (*t*) and a clustered TCR is assigned a score (*S*_match_) equal to the number of equivalent matches expected from a comparison between the query dataset and a shuffled background of size *N*_clust_. (a-b) Individual donor TCRs matching convergent clusters in the A3 and A5 GEX groups are shown in the red and yellow boxes along with annotations transferred from the matched clusters. (c) Bar plot showing the fraction of donor TCRs matched to convergent clusters, colored by GEX cluster group (same colors as in Fig. 2a). (d) Bar plot showing the fraction of donor TCRs matching to COVID19-associated clusters, colored by MHC restriction. (e) Matches to CDR3AA bias clusters are identified as statistically anomalous overlap between high-scoring regions for cluster-specific TCR and GEX scores. The TCR score (*S*_TCR_) measures the occurrence of enriched and depleted CDR3 amino acids, V genes, and CDR3 lengths for each cluster relative to background frequencies. Here *V* and *σ* are the V gene identifier and CDR3 amino acid sequence, respectively; *C_i_* is the CDR3AA cluster being matched; *L(σ)* is the length of the CDR3; and *σ^core^* represents the central portion of the CDR3, excluding the first four and last four residues. The GEX score (*S*_GEX_) measures expression of the top 50 differentially expressed genes (DEGs) specific to each CDR3AA cluster. Here *X* represents the standardized, log-transformed expression vector for the clonotype being scored, and DEG*_j_*is one of the differentially expressed genes for CDR3AA cluster C*_i_*with associated log2-fold change (log_2_FC) and Z-score (*Z*) from the differential expression calculation on the metaCoNGA dataset. (e-f) Panels are colored by TCR score (left), GEX score (middle), and combined score (right) for overlapping clonotypes that score well by both measures (overlap size and total number of clonotypes shown in panel heading along with *p-*value under null model where the scores are uncorrelated). (g-h) Fraction of total CD8+ and CD4+ clonotypes that match CDR3AA bias clusters, colored according to cluster functional group (**Fig. S10**).

To match a new single-cell dataset against the metaCoNGA CDR3AA bias clusters, we calculate cluster-specific TCR and GEX scores and look for clonotypes in the new dataset that score well by both metrics (**Fig. 6e-f** and Methods). The TCR sequence score for a given CDR3AA cluster (*S*_TCR_ in **Fig. 6e**) is defined by the enriched and depleted amino acid motifs, V genes, and CDR3 lengths of clonotypes in that cluster. The GEX score for a cluster (*S*_GEX_ in **Fig. 6e**) measures expression of the top differentially expressed genes (DEGs) in the metaCoNGA dataset for that cluster. This process is illustrated for individual donors from a recent study of colorectal cancer [32] matching to CD8+ CDR3AA cluster B6 (KIR+ CD8) and to CD4+ CDR3AA cluster D4 (tissue Tregs) in **Figure 6e-f**. Clonotypes with elevated *S*_TCR_ scores in their local GEX neighborhoods (red dots in the left panels) overlap significantly with clonotypes having elevated *S*_GEX_ scores (red dots in the middle panels). We define the specific clonotypes matching to each CDR3AA cluster (right panels) to be the subset of the overlap with the highest combined scores for both measures (see Methods). CD4+ and CD8+ CDR3AA cluster matches for all donors in this dataset are summarized by the bar plots in **Figure 6g-h**, in which matches to individual clusters have been condensed into functional groups for clarity (**Fig. S10**). From these plots we can see that there is statistically significant support for populations of T cells resembling the CDR3AA-biased clusters in these new samples, at combined frequencies from 5-10% of total clonotypes. Interestingly, the fractions of each dataset matching to individual clusters varies substantially from donor to donor. While this may be driven in part by technical issues such as sample size and noise in our matching procedure, we anticipate that these inferred population sizes carry information on immune state relevant to health and disease. For example, we found a statistically significant decline in the frequency of the CD8+ CDR3AA cluster B6 (KIR+ CD8) with increasing donor age in the metaCoNGA dataset, whereas the frequencies of the Treg CD4+ clusters tend to increase with age (**Fig. S11**).

## Discussion

To map connections between TCR sequence and cellular phenotype in an unbiased and comprehensive manner, we assembled a large collection of TCR sequences and associated gene expression profiles from 91 independent single-cell genomics studies spanning a range of contexts in human health and disease. The large size of the combined dataset is important for two reasons. First, it allows sensitive detection of statistically significant TCR sequence clusters (**Fig. 2**) despite the shallow depth of single-cell genomics profiling and the rarity of many clusters (a consequence of HLA restriction and variability in immune exposures). Second, it provides the statistical power to detect weak TCR sequence composition biases even without prior knowledge of the relevant cell populations or the nature of the sequence bias (**Figs. 3-5**). To leverage this massive dataset, we developed new algorithms for TCR sequence clustering and TCR-GEX covariation analysis, and a new feature-selection strategy for multimodal single-cell data on T cells that may generalize to other settings. Application of these new tools produced data resources that may have broad utility: a large collection of TCR sequence clusters and associated GEX profiles, many annotated with putative HLA restriction and/or predicted reactivity; a map of T cell populations with statistically significant TCR sequence composition biases that may reflect unique developmental, functional, and/or specificity features; a list of TCR-graph autocorrelated genes (TAGs) for use as variable gene features in single cell workflows that can help to alleviate batch effects while focusing clustering and dimensionality reduction on functionally relevant differences.

Beyond the dataset itself and these new analysis tools, we report a number of findings that may be of general interest. T cell clusters defined solely by TCR sequence similarity tend to have coherent gene expression patterns, and these expression patterns further subdivide based on the antigen source, with T cells specific for Flu, CMV, EBV, and SARS-CoV-2 mapping to distinct regions of the GEX landscape (**Fig. 2c**). In the case of SARS-CoV-2, some of this substructure may be explained by the presence of an ongoing COVID19 response at the time of sample collection: higher *MKI67* expression, indicative of proliferation, is visible in many of the SARS-CoV-2 clusters (**Fig. 2e**). Focusing on the largest TCR sequence cluster in each of the 21 gene expression groups, we were able to annotate a surprisingly large number with tentative MHC and/or antigen restriction (**Fig. 2b**), including many well-studied immunodominant responses. The gaps in these annotations should provide fruitful directions for future de-orphanization efforts as the responses likely correspond to undescribed highly immunodominant epitopes across the population.

At the other end of the TCR/GEX covariation spectrum, we identified multiple cell populations with significantly biased CDR3 sequence composition. The origins of these biases are varied: the greater hydrophobicity of regulatory T cells (**Fig. 5**) likely relates to their strong binding to self-pMHC, whereas the distinctive sequence biases of the thymocyte populations observed here (**Figs. 3+5**) are a consequence of their developmental stage (e.g., relative to positive and negative selection checkpoints). Our analysis identified multiple CD8+ populations with enrichment for large hydrophobic residues and elevated expression of marker genes including *ZNF683* (HOBIT), *IKZF2* (HELIOS) and multiple KIR genes. The size of the dataset and diversity of tissue and disease contexts enabled a finer-grained dissection of these cells into multiple subpopulations, some of which (e.g., B8 and B14 in **Figure 3**) may be distinct from the core HOBIT+/HELIOS+/KIR+ cells identified in previous work [4][25][24–27]. The presence of coherent TCR sequence biases is suggestive of unconventional developmental trajectories and/or antigen restriction for these CD8+ populations. A strength of this integrative analysis is that the populations it identifies need not be fully separated into traditional GEX clusters or subregions of the dimensionality-reduction landscape. Given the recent proliferation of studies describing variations of these unconventional cell types with shared phenotypic features (KIR+ etc.), this effort provides a categorization of potentially distinct ontogenic and functional diversity.

Our analysis has a number of limitations that could be addressed in future work. Combining single-cell gene expression data from different labs, tissues, and disease contexts can introduce batch effects (systematic differences between studies driven by experimental procedures and other technical artifacts) that confound detection of genuine biological differences. Our procedure for selecting variable genes using the TCR sequence neighbor graph (which has fewer technical biases) alleviates some of these effects (**Figs. S1-2**), but it is worth exploring more sophisticated batch correction strategies in the future. Relationships between TCR sequence and cellular phenotype are mediated by structural and biophysical properties of the TCR for which sequence is a weak proxy. As deep learning structural modeling methods continue to improve, it should become possible to identify TCR convergence and bias directly at the level of modeled TCR structures [33]. All the analyses in this work have been conducted at the level of TCR clonotypes, using gene expression profiles that are averaged over clonally-related cells. This allows us to identify relationships between TCR sequence and phenotype that transcend clonal boundaries, but it also masks biologically meaningful intra-clonotype phenotypic variation. Finally, it is important to recognize that by focusing on TCR-graph associated variable genes for GEX analyses, we may be missing orthogonal axes of T cell phenotypic variation.

## Methods

### Datasets and data processing

Single-cell datasets with GEX and TCR information for human T cells were identified through literature searches, GEO database (https://www.ncbi.nlm.nih.gov/geo/) queries, and from references on the 10X Genomics website (https://www.10xgenomics.com/publications?refinementList%5Bspecies%5D%5B0%5D=Human&refinementList%5BproductGroups%5D%5B0%5D=Universal%203%27%20Gene%20Expression). Gene expression counts information and TCR sequences were downloaded from multiple sources (the GEO database, author websites, the short read archive, etc) as indicated in **Table S1**. Where possible, GEX data was downloaded as sequence reads (.fastq files) and processed locally using cellranger (version 6.0.0 with default parameters). TCR sequence information was most often parsed from 10X Genomics output files (generally the filtered contigs CSV files), in which cases the ‘setup_10x_for_conga.py’ script in the CoNGA package [4] was used to identify spurious chain sharing (due to ambient TCR mRNA, cell doublets, and other technical artifacts) and define a cleaned set of TCR clonotypes. We only included studies which provided V and J gene identifiers as well as CDR3 amino acid sequences. Most studies also made CDR3 nucleotide sequences available, however for a few studies only the amino acid sequences could be downloaded. For those, we inferred the most parsimonious nucleotide sequence for each CDR3 amino acid sequence to minimize non-templated N-insertions.

GEX data was processed and merged using a python pipeline built with the scanpy software package [34]. The aligned GEX counts and TCR sequences for individual cells of each dataset were stored in a single AnnData [35] data structure. We excluded cells with fewer than 200 or more than 3500 expressed genes and cells with greater than 10% of their transcript counts coming from mitochondrial genes. Transcript counts were reduced to the common set of 11,430 genes shared across all of the studies, normalized to sum to 10000 within each cell, then log-transformed (x-->ln(x+1)) and averaged over the cells of each TCR clonotype. A TCR clonotype was defined as a unique, paired TCR sequence at the nucleotide level and occurring within a single study. Most studies also provided donor information which could have been used to subdivide clonotypes further since they should not be shared across donors, however, we saw many cases of high-frequency sharing of identical nucleotide sequences across donors suggesting noise in the donor assignments (for example due to noisy cell hashing assignments when running multiple samples in the same experiment). Artificially splitting exact-nucleotide clonotypes creates very strong signal of apparent but spurious convergent recombination that can skew the TCR clustering analysis, so we elected to use the more stringent, study-level clonotype definition, even if this occasionally led to merging of genuinely distinct T cell clonal lineages (for example in MAIT cells, where exact nucleotide sharing between subjects is possible in larger cohort studies).

Studies parsed from raw reads all contained a single donor per 10X Genomics sample and are labeled as such. Studies processed by the original authors often contained multiple donors per 10X Genomics sample distinguished by hashtag antibodies. In these cases the original hashtag annotations were used to assign cells to a donor. Tissue assignments per samples were done by manually cross-referencing the GSE series of each sample to GEO database. A small number of samples (0.4% of cells) contained multiple tissues that could not be resolved and were labeled as ‘mixed’. Assignment of the disease status for each sample was performed in a similar manner.

#### Defining CD4+ and CD8+ subsets

Most alpha-beta T cells are positive for one of either the CD4 or the CD8 co-receptors. This dichotomy determines a T cell’s MHC class restriction and biases its transcriptional profile. We and others have also observed [36] that CD4+ versus CD8+ status influences the sequence properties of the TCR, with CD4+ T cells having distinct V gene usage and CDR3 sequence composition (elevated charge, among other features) relative to CD8+ T cells. These sequence composition biases are modest at the level of individual TCRs and thus cannot be used to confidently assign CD4/CD8 status, but they are highly statistically significant when assessed at the level of large populations. Thus we elected to split the dataset into CD4+ and CD8+ subsets prior to CDR3 bias analysis (**Figs. 3-5**) in order to remove CD4 vs CD8 signal and define the most representative background sequence composition for each GEX neighborhood.

Unfortunately, defining CD4+ vs CD8+ status from gene expression alone is not straightforward: CD4 expression tends to be rather low, which together with generally high levels of dropout in single-cell data means that GEX-derived clusters and dimensionality reductions often blur the boundaries between CD4+ and CD8+ populations. Where present, per-cell counts for barcoded antibodies against surface-expressed CD4 and CD8 proteins can be used to refine these clusters, but only a subset of the studies included such antibodies in their workflows. Thus we took an iterative and fairly stringent approach to defining the CD4+ and CD8+ subsets, preferring to leave cells unclassified rather than include misclassified cells which could lead to “uninteresting” CDR3 amino acid biases and complicate interpretation.

In the first step, individual datasets split by study and by donor (where such annotation was available) were analyzed individually using the CoNGA pipeline, and clonotypes were assigned to CD4/CD8 on the basis of CD4 vs CD8 transcript counts for high-resolution Leiden clusters (’run_conga.py --subset_to_CD4_cells or --subset_to_CD8_cells’). The rationale for this first level of classification was our anecdotal observation that, when sufficient cells are present, datasets from individual donors gave the cleanest GEX clusters and dimensionality reductions (likely due to minimal noise from batch effects). Using high-resolution clusters as the units of classification is helpful because many individual cells have zero counts for all of CD4, CD8A, and CD8B (due to dropout, for example), yet they may be transcriptionally similar to other cells which do have nonzero counts for one or more of these coreceptor transcripts.

This per-donor approach appeared to work reasonably well when sufficient cells were present, but some donors contributed small numbers of cells to the dataset, and as noted above the donor assignments appeared noisy in some cases, so we applied an additional filter based on GEX clusters calculated over the entire merged dataset. Average CD4 and CD8 expression levels were plotted for each of the global GEX clusters, and a subset of high-confidence CD4+ and CD8+ clusters were identified. Membership in these clusters was used to filter (not expand) the CD4/CD8 assignments coming from the per-donor analysis: clonotypes initially assigned as CD4 (CD8) that belonged to a confident CD8+ (CD4+) global GEX cluster were excluded (set as unassigned). An initial round of CDR3 bias analysis suggested that some CD4+ clonotypes remained in the CD8 subset (and vice versa), and so we applied a final stringent filter based on transcript counts for the “wrong” coreceptor(s): clonotypes with a clone-averaged log-transformed CD4 transcript count greater than 0.2 were eliminated from the CD8 subset (and similarly for the CD4 subset). After application of this final filter, 2,599,503 of the 3,903,971 total clonotypes (66.6%) were assigned to the CD4+ subset, 815,535 (20.9%) were assigned to the CD8+ subset, and 488,933 (12.5%) were unassigned.

#### Identification of TCR neighbor graph-associated genes

To select variable genes for use as features in downstream GEX analyses (PCA, clustering, dimensionality reduction, GEX neighbor finding), we identified genes with statistically significant autocorrelation with the TCR neighbor graph (**Fig. S1b**). To assess graph autocorrelation, we used the hotspot Z score [16], a variant of Moran’s spatial autocorrelation index [37]. The TCR *K* nearest neighbor (KNN) graph was constructed by connecting each TCR clonotype to its 500 nearest neighbors as measured by TCRdist. The log-transformed normalized counts for each gene were standardized to have mean 0 and variance 1. The hotspot Z-score for a gene with standardized values *{g_i_}* is calculated by summing the product *g_i_*g_j_* over all edges *(i,j)* in the KNN graph *E*:

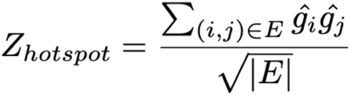

, where *|E|* equals the number of edges in the KNN graph *E*.

If the gene varies in a way that is uncorrelated with the TCR neighbor graph, then this sum will tend to be small and normally distributed. If instead the values of *g* for graph neighbors are correlated (i.e., knowing that *g_i_* is positive (or negative) increases the chance that *g_j_* is also positive (or negative) if *i* and *j* are graph neighbors), then the sum will be larger. The top 200 genes by hotspot Z score were selected, after excluding ribosomal and mitochondrial genes.

#### GEX clustering and dimensionality reduction

Python scripts built on the scanpy package were used for GEX data processing. As noted above, the raw transcript counts for the 11,430 genes common to all studies were normalized to sum to 10,000, then log+1 transformed and averaged over all cells in each TCR clonotype. The top 200 TCR-graph associated genes were selected as variable gene features. The clonotype-averaged, log-transformed, and normalized counts for these genes were standardized to have mean 0 and variance 1 with outlier values clipped at 10. The top 20 principal components were computed (scanpy.tl.pca), and nearest neighbors were identified using Euclidean distances in PC space (sc.pp.neighbors with n_neighbors=10) as input for Leiden clustering (scanpy.tl.leiden) and UMAP dimensionality reduction (scanpy.tl.umap).

#### Detection of statistically significant TCR sequence clusters

TCR sequence clusters were defined by a 2-step procedure. In the first step, TCR clonotypes with a greater than expected number of sequence neighbors were identified. In the second step, these neighbor-enriched clonotypes were grouped into clusters of similar sequences. To identify significantly neighbor-enriched TCR sequences, a shuffled set of background sequences was first constructed by splitting and remixing the V(D)J recombinations seen in the observed (foreground) sequences. This shuffling was performed at the nucleotide level, and V(D)J recombination scenarios were assigned to foreground sequences by parsimony to minimize the number of non-templated N nucleotides. The reshuffled sequences were filtered to remove out-of-frame sequences and stop codons and to match the length and N-insertion distribution of the observed sequences. The V and J gene usage distributions of the shuffled sequences matched the gene usage distribution of the observed sequences, and for the TCR alpha chain the V-J gene pairing distributions were also constrained to match, since there are significant biases in Va-Ja gene pairing that arise from the fact that failed recombination events can be rescued by subsequent recombinations on the same DNA chain. This shuffling procedure was performed independently for the alpha and beta chains, yielding background TCRa and TCRb sequences. To estimate the likelihood of seeing an observed number of sequence neighbors for a given TCR sequence, background TCRdist distributions were calculated independently for the TCRa sequence and for the TCRb sequence. A paired background TCRdist distribution was then constructed from these single-chain distance distributions using the assumption of random a/b chain pairing and the fact that the paired TCRdist equals the sum of the alpha-chain and beta-chain distances. Under these assumptions, the paired TCRdist distribution from matching to a background set consisting of all shuffled TCRa-TCRb combinations is given by the convolution of the single-chain distributions. This allows us to calculated the expected frequency distribution for *N^2^*paired TCRdist distances using only the *2*N* single-chain distances. For the results reported here, we used single-chain background sets consisting of *N=*250,000 sequences, corresponding to an effective paired background set of size 6.25e10, which is about 20,000 times larger than the foreground set of 3 million clonotypes. This expected background TCRdist distance distribution was calculated for each observed TCR clonotype, and the probability of seeing an observed number of foreground neighbors with distance less than or equal to a given threshold T was estimated using the Poisson distribution. Here we used four TCRdist distance thresholds: T = 24, 48, 72, and 96. To account for multiple testing, the neighbor-count probabilities were multiplied by 4*C where C was the number of observed foreground clonotype sequences (C=3,971,194).

TCR sequences with a statistically significant number of neighbors at one or more of the 4 TCRdist distance thresholds were grouped into clusters using a simple greedy graph-based clustering algorithm. A directed neighbor graph was constructed by looping over the four distance thresholds T in {24,48,72,96} and adding an edge from TCRi to TCRj if TCRi has a significantly-enriched neighbor number at threshold T and TCRdist(TCRi, TCRj)<=T. TCR sequence clusters were formed from this graph by repeatedly selecting the node with the highest number of outgoing edges as a new cluster center, assigning all of its connected neighbors as cluster members, and deleting the cluster center and all cluster members from the graph. We chose this strategy after exploring a variety of distance and graph-based clustering methods mainly for its simplicity and because it bounds the maximal sequence distance within clusters: all cluster members are guaranteed to be within a distance threshold T of the cluster center. This prevents clusters from spreading too far across sequence space and increases the likelihood of shared reactivity, with the downside being that single specificities may be split across multiple sequence clusters (for example, MAIT cell TCRs with TRBV20 chains cluster separately from MAIT cells with TRBV6-family chains).

#### Annotations of TCR sequence clusters

Potential epitope specificity was assigned to individual TCR sequences by matching to a filtered version of the VDJdb [38] database. Matches were identified using the TCRdist measure, with statistical significance assigned by matching each TCR to a large shuffled background database to create an expected probability distribution of TCRdist scores. HLA and CMV association *p*-values for each convergent TCR cluster were calculated by tracking occurrences of that cluster’s TCR beta chains in the ∼650 repertoires of the Emerson CMV cohort [18], for which CMV serostatus and HLA genotyping is available. The significance of an observed overlap between a given cluster’s occurrence pattern and an HLA allele (or CMV seropositivity) was assessed with the hypergeometric distribution.

#### Identification of CDR3 amino acid sequence biases in GEX neighborhoods

To identify regions in GEX space with biased CDR3 amino acid sequences, we analyzed the TCR sequence distributions in neighborhoods (a central clonotype and its connected neighbors) of the GEX KNN graph with K=5000 neighbors. To construct the GEX graph, we first normalized the raw transcript counts for each cell to sum to 10,000. These normalized counts were log transformed (x-->ln(x+1)) and averaged over the cells in each T cell clonotype. The top 200 TCR-associated genes (**Fig. S1b**) were selected as variable gene features for GEX analysis using a pipeline implemented in the scanpy [34] python package. The log-normalized counts were standardized to have mean 0 and variance 1 and clipped to remove values larger than 10. The top 20 principal components were computed, and the KNN graph was built using Euclidean distances in this 20-dimensional PC space.

For each GEX neighborhood consisting of a central clonotype and its K=5000 nearest neighbors, we computed the frequencies of each the 20 amino acids in the 5001 CDR3alpha sequences and the 5001 CDR3beta sequences. The central portion of each CDR3 sequence was analyzed, excluding the first 4 and last 4 amino acids which are less likely to directly contact the pMHC. The observed CDR3a (CDR3b) frequency distribution for each neighborhood was compared to the background distribution calculated from the full set of CDR3a (CDR3b) sequences, and a chi-squared distribution with 19 degrees of freedom was used to assign a *P* value to the observed divergence. These *P* values were multiplied by the total number of neighborhoods (i.e., clonotypes) to yield *E* values that account for multiple testing, and clonotypes with a CDR3a or CDR3b *E* value less than a threshold of 1e-6 were selected for further analysis (**Figs. 3a** and **5a**). Because CD4 and CD8 T cells are known to have distinct TCR sequence biases, this analysis was performed on the CD4+ and CD8+ subsets of the dataset independently. In addition, the large cluster of CD8+ MAIT cell clonotypes (the light blue cluster in **Fig. S1c**) was excluded from this analysis so as not to skew the background CDR3 sequences or swamp weaker signals. To create a landscape of CDR3 sequence bias, each significant neighborhood (*E* value<1e-6) was mapped to a 40-dimensional vector consisting of the 20 CDR3a AA frequencies followed by the 20 CDR3b AA frequencies. Clustering and dimensionality reduction were applied to these vectors using a workflow implemented in scanpy: the AA frequency vectors were first standardized to have mean 0 and variance 1 in each dimension, then the UMAP dimensionality reduction and Leiden clustering methods were applied to generate 2D landscapes and cluster assignments (**Figs. 3c and 5c**).

#### Annotating new datasets with metaCoNGA features

Two search algorithms can be used to annotate a new single-cell dataset using the metaCoNGA features. First, statistically significant matches between TCR sequences in the new dataset and metaconga TCR clusters are identified by using the TCRdist metric to match each sequence to a database of representative TCRs from metaconga TCR clusters. Rather than matching against all 109,384 significantly neighbor-enriched TCRs, we focus on the 2173 convergent clusters with at least 10 members, downsampling each cluster to 10 TCRs (the central TCR in the cluster and 9 additional randomly selected TCRs). This reduces the nominal multiple testing burden (which equals the product of the number of foreground TCRs and the number of database TCRs) and equalizes matching across metaconga clusters of different sizes. To assess the significance of a match between a foreground TCR and a metaconga TCR, the observed TCRdist distance is compared to a background distribution generated for that specific foreground TCR by matching to a shuffled set of background TCRs. The background TCR set is created from the foreground set as described above in the section on TCR convergence detection. Note that the background TCRdist distribution is TCR-specific: a foreground TCR with shorter CDR3 regions that are closer to germline will tend to have an excess of smaller TCRdist distances compared with one that has long CDR3s.

Potential matches to the CDR3AA bias clusters are identified by looking for regions that match both the TCR and GEX features of each cluster (**Fig. 6**). The dataset is first partitioned into CD4+ and CD8+ subsets which are matched against the CD4+ and CD8+ CDR3AA clusters, respectively. A TCR score (*S*_TCR_) is defined based on enrichment of CDR3 amino acids, V genes, and CDR3 lengths for each cluster relative to background frequencies:

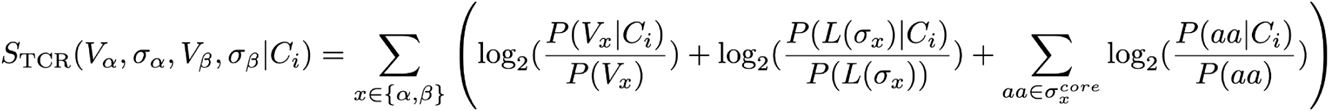

Here *V* and *σ* are the V gene identifier and CDR3 amino acid sequence, respectively; *C_i_* is the AA-cluster being matched; *L(σ)* is the length of the CDR3; and *σ^core^* represents the central portion of the CDR3, excluding the first four and last four residues. The TCR score is averaged over local GEX neighborhoods in the query dataset to reduce noise and better capture the statistical nature of the amino acid sequence biases (KNN neighborhoods with K= 2.5% of the number of clonotypes).

A GEX score (*S*_GEX_) is defined based on expression of the top 50 differentially expressed genes (DEGs) specific to each CDR3AA cluster:

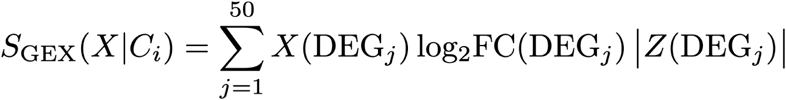

Here *X* represents the standardized, log-transformed expression vector for the clonotype being scored, and DEG*_j_* is one of the differentially expressed genes for CDR3AA cluster C*_i_* with associated log2-fold change (log_2_FC) and Z-score (*Z*) from the differential expression calculation on the metaCoNGA dataset.

For each AA-cluster, the top-scoring regions for *S*_TCR_ and *S*_GEX_ are identified and their overlap is calculated. Since we don’t know in advance the expected frequency of the matching population, a range of top-scoring fractions (10%, 5%, 2.5%, and 1% of the dataset) are evaluated. For each fraction, the top-scoring subsets of that size by *S*_TCR_ and *S*_GEX_ are identified and the statistical significance of their overlap is measure using the hypergeometric distribution. The most significant overlap that exceeds a specified *p-*value threshold (here 10^−3^) is used to define the set of clonotypes that match the given cluster. The number of matching clonotypes is taken to be the difference between the observed and expected overlap sizes (ie, the excess number of overlapping clonotypes), with the specific clonotypes within the overlapping region chosen on the basis of their combined TCR and GEX scores (individually rank-transformed and then summed). Matches are first identified for each CDR3AA cluster individually; then clonotypes matching multiple clusters are assigned to the cluster with the highest combined TCR and GEX score. Here, multiple matches can occur because of similarities between the sequence and phenotypic features of distinct CDR3AA clusters (between different tissue Treg clusters, for example, or the HOBIT/HELIOS/KIR+ clusters).

## Data Availability

Processed datasets, convergent clusters, and related resources will be made available prior to final publication, subject to patient privacy constraints. Check https://github.com/phbradley/metaconga for updates.

## Code Availability

Analysis software will be made available prior to final publication. Check https://github.com/phbradley/metaconga for updates.

## Acknowledgments

This research was made possible by the efforts of many researchers who have diligently generated, annotated, and shared their datasets with the broader scientific community. We extend our sincere thanks for their contributions, noting particular assistance from Jim Heath, Yapeng Su, Sarah Teichmann, John Tsang, and Zemin Zhang in accessing and parsing their datasets. This work was supported through NIAID grants R01AI136514 (P.G.T and P.B.), U01AI150747 (P.G.T), R21AI169085 (P.G.T and P.B.), and T32AR007258 (K.S.), and NIGMS grant R35GM141457 (P.B.). Scientific Computing Infrastructure at Fred Hutch was funded by NIH ORIP grant S10OD028685. American Lebanese Syrian Associated Charities at St. Jude (P.G.T). A.-C.V. acknowledges funding support from the COVID-19 Clinical Trials Pilot grant from the Executive Committee on Research at MGH, a COVID-19 Chan Zuckerberg Initiative grant (2020-216954), the funds from the Manton Foundation and the Klarman Family Foundation, and the National Institutes of Health (DP2CA247831).

## MGH COVID-19 Collection & Processing Team

<u>Leadership Team</u>

Michael Filbin

Department of Emergency Medicine, Massachusetts General Hospital, Boston, MA 02115

Nir Hacohen, Moshe Sade-Feldman

Massachusetts General Hospital Cancer Center, Boston, MA 02115

Marcia Goldberg, Roby Bhattacharyya

Division of Infectious Diseases, Department of Medicine, Massachusetts General Hospital, Boston, MA 02115

<u>Collection Team</u>

Kendall Lavin-Parsons, Blair Parry, Brendan Lilley, Carl Lodenstein, Brenna McKaig, Nicole Charland, Hargun Khanna, Justin Margolin

Department of Emergency Medicine, Massachusetts General Hospital, Boston, MA 02115

<u>Processing Team</u>

Moshe Sade-Feldman, Anna Gonye, Irena Gushterova, Tom Lasalle, Nihaarika Sharma Massachusetts General Hospital Cancer Center, Boston, MA 02115

Brian C. Russo, Maricarmen Rojas-Lopez

Division of Infectious Diseases, Department of Medicine, Massachusetts General Hospital, Boston, MA 02115

Kasidet Manakongtreecheep, Jessica Tantivit, Molly Fisher Thomas, Thomas Lasalle, Thomas Eisenhaure Massachusetts General Hospital Center for Immunology and Inflammatory Diseases, Boston, MA 02115

<u>Single-cell genomics team</u>

Pritha Sen, Christopher Cosgriff, Jessica Tantivit, Tom Lasalle, Thomas Eisenhaure, Kasidet Manakongtreecheep, Benjamin Arnold, Alice Tirard, Miguel Reyes, Rachelly Normand, Steven Blum, Swetha Ramesh

Massachusetts General Hospital Center for Immunology and Inflammatory Diseases, Boston, MA 02115

## Author Contributions

S.S, P.G.T, and P.B. conceptualized the study. P.G.T and P.B. acquired funding and supervised the study. S.S. and P.B. conducted the investigation and formal analysis, and generated visualizations. S.S., K.V., W.D.H., and P.B. identified and processed the publicly available single-cell datasets and cleaned the accompanying metadata. S.S., S.V., and P.B. developed the methodology and software used for the study. K.S., N.S., MGH COVID-19 Team, and A.-C.V. provided datasets for the study. S.S, K.V., P.G.T, and P.B wrote the original draft of the mansucript. S.S, A.M., P.G.T, and P.B performed extensive review and editing of the original manuscript.

## Supplementary Tables

**Table S1. Source datasets and literature citations**

**Table S2. Top 2000 TCR-graph autocorrelated genes (TAGs).** Note that only the top 200 were used for GEX analyses.

**Table S3. Differentially-expressed genes for the 21 GEX groups of convergent TCR clusters (A0-A20)**

## Supplementary Figures

**Figure S1. Identification and application of universal features of human T cells provide robust single-cell GEX integration.**

**Figure S2. Comparison of dimensionality reduction and clustering results using HVGs or TAGs**.

**Figure S3. Pipeline for TCR convergence analysis.**

**Figure S4. Convergent TCR clusters have coherent gene expression profiles.**

**Figure S5. Consistency of CD4 versus CD8 gene expression for convergent TCR clusters.**

**Figure S6. Pipeline for CDR3AA bias analysis.**

**Figure S7. CDR3AA bias landscapes colored by additional TCR sequence features.**

**Figure S8. Top differentially expressed genes for the CD8+ CDR3AA bias clusters.**

**Figure S9. Top differentially expressed genes for the CD4+ CDR3AA bias clusters.**

**Figure S10. Putative functional groupings of the CD8+ and CD4+ CDR3AA bias clusters as used in the bar plots in main text Figure 6**.

**Figure S11. CDR3AA-bias cluster matches as a function of donor age in the metaCoNGA dataset.**

## References

1. Pellicci DG, Koay H-F, Berzins SP. Thymic development of unconventional T cells: how NKT cells, MAIT cells and γδ T cells emerge. Nat Rev Immunol. 2020;20: 756–770. doi:10.1038/s41577-020-0345-y

2. Dash P, Fiore-Gartland AJ, Hertz T, Wang GC, Sharma S, Souquette A, et al. Quantifiable predictive features define epitope-specific T cell receptor repertoires. Nature. 2017;547: 89–93. doi:10.1038/nature22383

3. Glanville J, Huang H, Nau A, Hatton O, Wagar LE, Rubelt F, et al. Identifying specificity groups in the T cell receptor repertoire. Nature. 2017; 1–17. doi:10.1038/nature22976

4. Schattgen SA, Guion K, Crawford JC, Souquette A, Barrio AM, Stubbington MJT, et al. Integrating T cell receptor sequences and transcriptional profiles by clonotype neighbor graph analysis (CoNGA). Nat Biotechnol. 2022;40: 54–63. doi:10.1038/s41587-021-00989-2

5. Schmidt F, Fields HF, Purwanti Y, Milojkovic A, Salim S, Wu KX, et al. In-depth analysis of human virus-specific CD8+ T cells delineates unique phenotypic signatures for T cell specificity prediction. Cell Rep. 2023;42: 113250. doi:10.1016/j.celrep.2023.113250

6. Chen DG, Xie J, Su Y, Heath JR. T cell receptor sequences are the dominant factor contributing to the phenotype of CD8+ T cells with specificities against immunogenic viral antigens. Cell Rep. 2023;42: 113279. doi:10.1016/j.celrep.2023.113279

7. Xie J, Chen DG, Chour W, Ng RH, Zhang R, Yuan D, et al. APMAT analysis reveals the association between CD8 T cell receptors, cognate antigen, and T cell phenotype and persistence. Nat Commun. 2025;16: 1402. doi:10.1038/s41467-025-56659-3

8. Newell EW, Sigal N, Nair N, Kidd BA, Greenberg HB, Davis MM. Combinatorial tetramer staining and mass cytometry analysis facilitate T-cell epitope mapping and characterization. Nat Biotechnol. 2013;31: 623–629. doi:10.1038/nbt.2593

9. Lagattuta KA, Kang JB, Nathan A, Pauken KE, Jonsson AH, Rao DA, et al. Repertoire analyses reveal T cell antigen receptor sequence features that influence T cell fate. Nat Immunol. 2022. doi:10.1038/s41590-022-01129-x

10. Textor J, Buytenhuijs F, Rogers D, Gauthier ÈM, Sultan S, Wortel IMN, et al. Machine learning analysis of the T cell receptor repertoire identifies sequence features of self-reactivity. Cell Syst. 2023;14: 1059–1073.e5. doi:10.1016/j.cels.2023.11.004

11. Lagattuta KA, Kohlgruber AC, Abdelfattah NS, Nathan A, Rumker L, Birnbaum ME, et al. The T cell receptor sequence influences the likelihood of T cell memory formation. Cell Rep. 2025;44: 115098. doi:10.1016/j.celrep.2024.115098

12. Park J-E, Botting RA, Conde CD, Popescu D-M, Lavaert M, Kunz DJ, et al. A cell atlas of human thymic development defines T cell repertoire formation. Science. 2020;367. doi:10.1126/science.aay3224

13. Terekhova M, Swain A, Bohacova P, Aladyeva E, Arthur L, Laha A, et al. Single-cell atlas of healthy human blood unveils age-related loss of NKG2C+GZMB-CD8+ memory T cells and accumulation of type 2 memory T cells. Immunity. 2024;57: 188–192. doi:10.1016/j.immuni.2023.12.014

14. Liu C, Martins AJ, Lau WW, Rachmaninoff N, Chen J, Imberti L, et al. Time-resolved systems immunology reveals a late juncture linked to fatal COVID-19. Cell. 2021;184: 1836–1857.e22. doi:10.1016/j.cell.2021.02.018

15. COvid-19 Multi-omics Blood ATlas (COMBAT) Consortium. Electronic address, COvid-19 Multi-omics Blood ATlas (COMBAT) Consortium. A blood atlas of COVID-19 defines hallmarks of disease severity and specificity. Cell. 2022;185: 916–938.e58. doi:10.1016/j.cell.2022.01.012

16. DeTomaso D, Yosef N. Hotspot identifies informative gene modules across modalities of single-cell genomics. Cell Syst. 2021;12: 446–456.e9. doi:10.1016/j.cels.2021.04.005

17. Pogorelyy MV, Minervina AA, Shugay M, Chudakov DM, Lebedev YB, Mora T, et al. Detecting T cell receptors involved in immune responses from single repertoire snapshots. PLoS Biol. 2019;17: e3000314. doi:10.1371/journal.pbio.3000314

18. Emerson RO, DeWitt WS, Vignali M, Gravley J, Hu JK, Osborne EJ, et al. Immunosequencing identifies signatures of cytomegalovirus exposure history and HLA-mediated effects on the T cell repertoire. Nat Genet. 2017;49: 659–665. doi:10.1038/ng.3822

19. Schumacher TN, Thommen DS. Tertiary lymphoid structures in cancer. Science. 2022;375: eabf9419. doi:10.1126/science.abf9419

20. Verstichel G, Vermijlen D, Martens L, Goetgeluk G, Brouwer M, Thiault N, et al. The checkpoint for agonist selection precedes conventional selection in human thymus. Sci Immunol. 2017;2. doi:10.1126/sciimmunol.aah4232

21. Malhotra N, Qi Y, Spidale NA, Frascoli M, Miu B, Cho O, et al. SOX4 controls invariant NKT cell differentiation by tuning TCR signaling. J Exp Med. 2018;215: 2887–2900. doi:10.1084/jem.20172021

22. Billiet L, De Cock L, Sanchez GS, Mayer RL, Goetgeluk G, De Munter S, et al. Single-cell profiling identifies a spectrum of human unconventional intraepithelial T lineage cells. bioRxiv. 2022. p. 2022.05.24.492634. doi:10.1101/2022.05.24.492634

23. Uehara S, Grinberg A, Farber JM, Love PE. A role for CCR9 in T lymphocyte development and migration. J Immunol. 2002;168: 2811–2819. doi:10.4049/jimmunol.168.6.2811

24. Billiet L, De Cock L, Sanchez Sanchez G, Mayer RL, Goetgeluk G, De Munter S, et al. Single-cell profiling identifies a novel human polyclonal unconventional T cell lineage. J Exp Med. 2023;220. doi:10.1084/jem.20220942

25. Li J, Zaslavsky M, Su Y, Guo J, Sikora MJ, van Unen V, et al. KIR+CD8+ T cells suppress pathogenic T cells and are active in autoimmune diseases and COVID-19. Science. 2022;376: eabi9591. doi:10.1126/science.abi9591

26. Choi SJ, Koh J-Y, Rha M-S, Seo I-H, Lee H, Jeong S, et al. KIR+CD8+ and NKG2A+CD8+ T cells are distinct innate-like populations in humans. Cell Rep. 2023;42: 112236. doi:10.1016/j.celrep.2023.112236

27. Lu BY, Lucca LE, Lewis W, Wang J, Nogueira CV, Heer S, et al. Circulating tumor-reactive KIR+CD8+ T cells suppress anti-tumor immunity in patients with melanoma. Nat Immunol. 2025;26: 82–91. doi:10.1038/s41590-024-02023-4

28. Chen X, Ghanizada M, Mallajosyula V, Sola E, Capasso R, Kathuria KR, et al. Differential roles of human CD4+ and CD8+ regulatory T cells in controlling self-reactive immune responses. Nat Immunol. 2025;26: 230–239. doi:10.1038/s41590-024-02062-x

29. Loh L, Carcy S, Krovi HS, Domenico J, Spengler A, Lin Y, et al. Unraveling the phenotypic states of human innate-like T cells: Comparative insights with conventional T cells and mouse models. Cell Rep. 2024;43: 114705. doi:10.1016/j.celrep.2024.114705

30. Chou C, Zhang X, Krishna C, Nixon BG, Dadi S, Capistrano KJ, et al. Programme of self-reactive innate-like T cell-mediated cancer immunity. Nature. 2022;605: 139–145. doi:10.1038/s41586-022-04632-1

31. Oh DS, Kim E, Normand R, Lu G, Shook LL, Lyall A, et al. SARS-CoV-2 infection elucidates features of pregnancy-specific immunity. Cell Rep. 2024;43: 114933. doi:10.1016/j.celrep.2024.114933

32. Chen Y, Wang D, Li Y, Qi L, Si W, Bo Y, et al. Spatiotemporal single-cell analysis decodes cellular dynamics underlying different responses to immunotherapy in colorectal cancer. Cancer Cell. 2024;42: 1268–1285.e7. doi:10.1016/j.ccell.2024.06.009

33. Bradley P. Structure-based prediction of T cell receptor:peptide-MHC interactions. Elife. 2023;12. doi:10.7554/eLife.82813

34. Wolf FA, Angerer P, Theis FJ. SCANPY: large-scale single-cell gene expression data analysis. Genome Biol. 2018;19: 15. doi:10.1186/s13059-017-1382-0

35. Virshup I, Rybakov S, Theis FJ, Angerer P, Wolf FA. anndata: Access and store annotated data matrices. J Open Source Softw. 2024;9: 4371. doi:10.21105/joss.04371

36. Emerson R, Sherwood A, Desmarais C, Malhotra S, Phippard D, Robins H. Estimating the ratio of CD4+ to CD8+ T cells using high-throughput sequence data. J Immunol Methods. 2013;391: 14–21. doi:10.1016/j.jim.2013.02.002

37. Moran PAP. Notes on continuous stochastic phenomena. Biometrika. 1950;37: 17–23. doi:10.1093/biomet/37.1-2.17

38. Shugay M, Bagaev DV, Zvyagin IV, Vroomans RM, Crawford JC, Dolton G, et al. VDJdb: a curated database of T-cell receptor sequences with known antigen specificity. Nucleic Acids Res. 2018;46: D419–D427. doi:10.1093/nar/gkx760

