## Supplementary Data for "Diverse modes of T cell receptor sequence convergence define unique functional and cellular phenotypes"

### Supplementary Tables

**Table S1. Source datasets and literature citations**

**Table S2. Top 2000 TCR-graph autocorrelated genes (TAGs).** Note that only the top 200 were used for GEX analyses.

**Table S3. Differentially-expressed genes for the 21 GEX groups of convergent TCR clusters (A0-A20)**

### Supplementary Figures

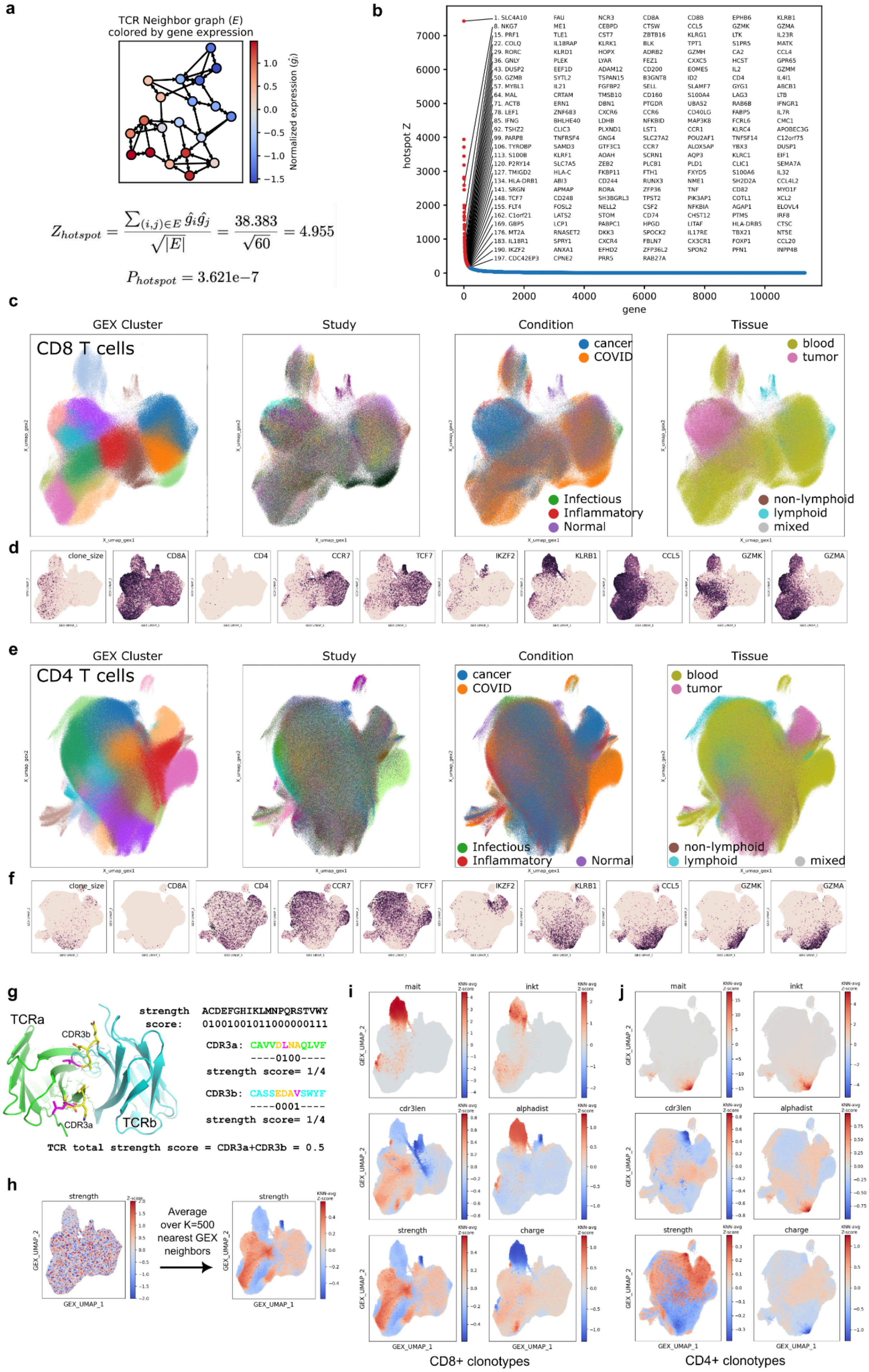
a
TCR Neighbor graph (E)
colored by gene expression
Normalized expression ($\hat{g}_i$)
$Z_{hotspot} = \frac{\sum_{(i,j)\in E} \hat{g}_i \hat{g}_j}{\sqrt{|E|}} = \frac{38.383}{\sqrt{60}} = 4.955$
$P_{hotspot} = 3.621e{-7}$
b
hotspot Z
gene
c
GEX Cluster
Study
Condition
Tissue
CD8 T cells
cancer
COVID
Infectious
Inflammatory
Normal
blood
tumor
non-lymphoid
lymphoid
mixed
d
clone_size
CD8A
CD4
CCR7
TCF7
IKZF2
KLRB1
CCL5
GZMK
GZMA
e
GEX Cluster
Study
Condition
Tissue
CD4 T cells
cancer
COVID
Infectious
Inflammatory
Normal
blood
tumor
non-lymphoid
lymphoid
mixed
f
clone_size
CD8A
CD4
CCR7
TCF7
IKZF2
KLRB1
CCL5
GZMK
GZMA
g
TCRa
CDR3b
CDR3a
TCRb
strength score: ACDEFGHIKLMNPQRSTVWY
01001001011000000111
CDR3a: CAVVDLNAQLVF
----0100----
strength score= 1/4
CDR3b: CASSEDAVSWYF
----0001----
strength score= 1/4
TCR total strength score = CDR3a+CDR3b = 0.5
h
strength
Average over K=500 nearest GEX neighbors
GEX_UMAP_1
GEX_UMAP_2
i
mait
inkt
cdr3len
alphadist
strength
charge
KNN-avg Z-score
CD8+ clonotypes
j
mait
inkt
cdr3len
alphadist
strength
charge
CD4+ clonotypes

**Figure S1. Identification and application of universal features of human T cells provide robust single-cell GEX integration. (**a) Toy example showing a K=3 nearest neighbors TCR graph colored by normalized expression of a gene whose expression is correlated with graph structure, in the sense that neighbors in the graph tend to have more similar expression than expected by chance. This graph correlation is captured by the hotspot Z score as shown in the equation below the graph. (b) Scatter plot of the TCR-graph autocorrelation Z-score (y-axis) versus gene rank (x-axis), with the top 200 TCR-graph autocorrelated genes (TAGs) shown in red and labeled. (c) 2D GEX landscape of CD8+ T cell clonotypes colored by GEX cluster (left), study (middle left), disease condition (middle right), and tissue (right). (d) 2D GEX landscape of CD8+ T cell clonotypes colored by (left to right) clone size, and expression of *CD8A*, *CD4*, *CCR7*, *TCF7*, *IKZF2*, *KLRB1*, *CCL5*, *GZMK*, and *GZMA*. (e) 2D GEX landscape of CD4+ T cell clonotypes colored by GEX cluster (left), study (middle left), disease condition (middle right), and tissue (right). (f) 2D GEX landscape of CD4+ T cell clonotypes colored by (left to right) clone size, and expression of *CD8A*, *CD4*, *CCR7*, *TCF7*, *IKZF2*, *KLRB1*, *CCL5*, *GZMK*, and *GZMA*. (g) An example showing the calculation of a CDR3 sequence feature (the 'strength' score, which counts the number of amino acids capable of forming strong interactions with pMHC) for a single TCR (3D view from the pMHC's perspective on the left). The per-position score is averaged over the central window (excluding the first and last four residues) of the CDR3 and summed across the two chains. (h) A plot of the raw score on the GEX landscape appears random, but averaging the score over GEX neighborhoods (each clonotype and its K=500 nearest GEX neighbors) reveals regions of the GEX landscape with high and low average score. (i) Six TCR sequence scores averaged over CD8+ GEX neighborhoods and mapped onto the 2D GEX landscape. (j) Six TCR sequence scores averaged over CD4+ GEX neighborhoods and mapped onto the 2D GEX landscape.

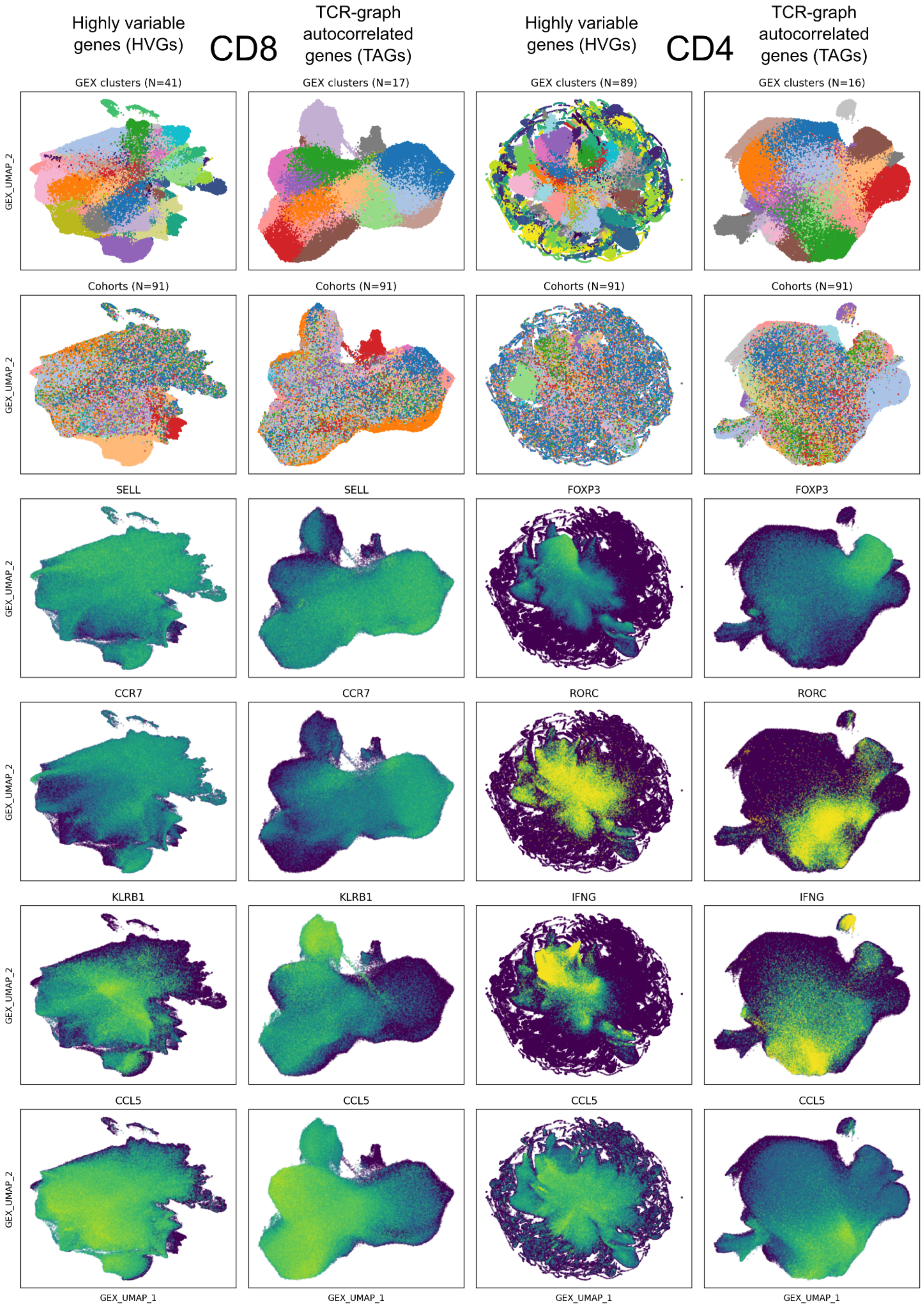

Highly variable genes (HVGs)
CD8
TCR-graph autocorrelated genes (TAGs)
Highly variable genes (HVGs)
CD4
TCR-graph autocorrelated genes (TAGs)
GEX clusters (N=41)
GEX clusters (N=17)
GEX clusters (N=89)
GEX clusters (N=16)
Cohorts (N=91)
SELL
FOXP3
CCR7
RORC
KLRB1
IFNG
CCL5
GEX_UMAP_1
GEX_UMAP_2

**Figure S2. Comparison of dimensionality reduction and clustering results using HVGs or TAGs**. The CD8+ (left two columns) and CD4+ (right two columns) datasets were processed using as features either the top 200 highly variable genes (HVGs, columns 1 and 3) or the top 200 TCR-graph autocorrelated genes (TAGs, columns 2 and 4). The top 20 GEX PCs were used for K=10 nearest neighbor finding. Leiden clustering was performed with resolution parameter=1.0, resulting in the indicated number of clusters in each case (first row). Default UMAP projections were generated from the KNN graphs. Raw transcript counts for the indicated genes were standardized to sum to 10,000 within each cell, then log+1-transformed and normalized to have mean 0.0 and standard deviation of 1.0 (clipping values at +/- 10), and plotted on the UMAP landscapes. Each dot corresponds to a single clonotype, sorted by increasing gene expression, with an alpha factor of 0.2 to reduce overplotting. Examination of the clustering and dimensionality results suggested to us that the TAG-based processing yielded improved results, with fewer and more interpretable clusters and clearer differentiation of subsets such as MAIT cells (KLRB1-high), naive cells (CCR7- and SELL-high), and regulatory T cells (FOXP3-high).

### Pipeline for TCR convergence analysis (main Fig. 2)

#### Step 1
**Identify TCR neighbors**

- Compute all-vs-all TCRdist distance matrix.
- For each TCR, identify nearby TCRs at four distance thresholds: 24, 48, 72, and 96

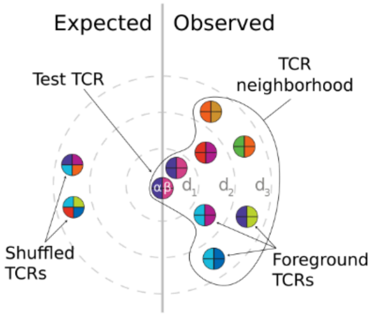


#### Step 2
**Compute neighbor-enrichment *P* values**

- Construct background TCR set by VDJ shuffling
- For each TCR, count neighbors in background
- Assign neighbor-enrichment P values by comparing observed to expected neighbor #s

#### Step 3
**Cluster neighbor-enriched TCRs**

- Build a directed graph of all significantly neighbor-enriched TCRs, connecting ti to tj if TCRdist(ti,tj)<T and ti is significant at threshold T.
- Select TCR with most neighbors, make cluster 0 from that TCR and its graph neighbors, then delete those nodes from the graph and find the TCR with the most neighbors, repeat

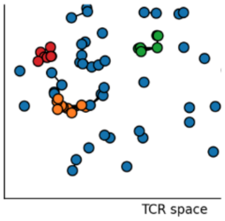


#### Step 4
**Visualize TCR clusters**

- Construct paired V/J/CDR3 sequence logos.
- Map TCRs back to GEX and metadata to get average gene expression, tissue composition, and cohort occurrence for each cluster.

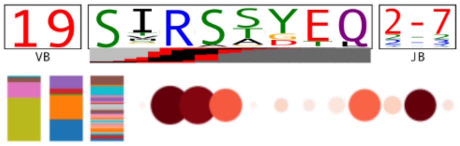


#### Step 5
**Construct GEX landscape of TCR clusters**

- Run PCA using as features the cluster-averaged gene expression values for the top 200 TCR-hotspot genes
- Run UMAP on the cluster PCs to create a 2D lanscape
- Run Leiden clustering on the cluster PCs to define GEX groups

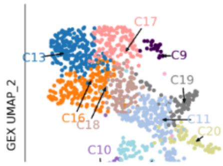


#### Step 6
**Compute cluster HLA and CMV associations**

- For each cluster, look for occurrences of cluster beta chains in HLA- and CMV-typed bulk repertoire cohort
- Assess statistical correlation between cluster occurrence pattern and occurrence patterns for each HLA allele, and for CMV seropositivity

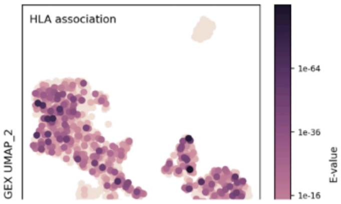


#### Step 7
**Match clusters to literature TCR databases**

- Match cluster TCRs to antigen-specific TCRs from published datasets using TCRdist
- Assess significance of matches using comparisons to a VDJ-shuffled background

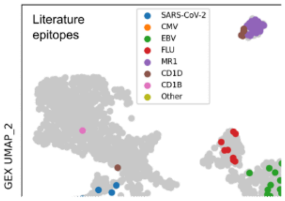

**Figure S3. Pipeline for TCR convergence analysis.** Details can be found in the Methods section.

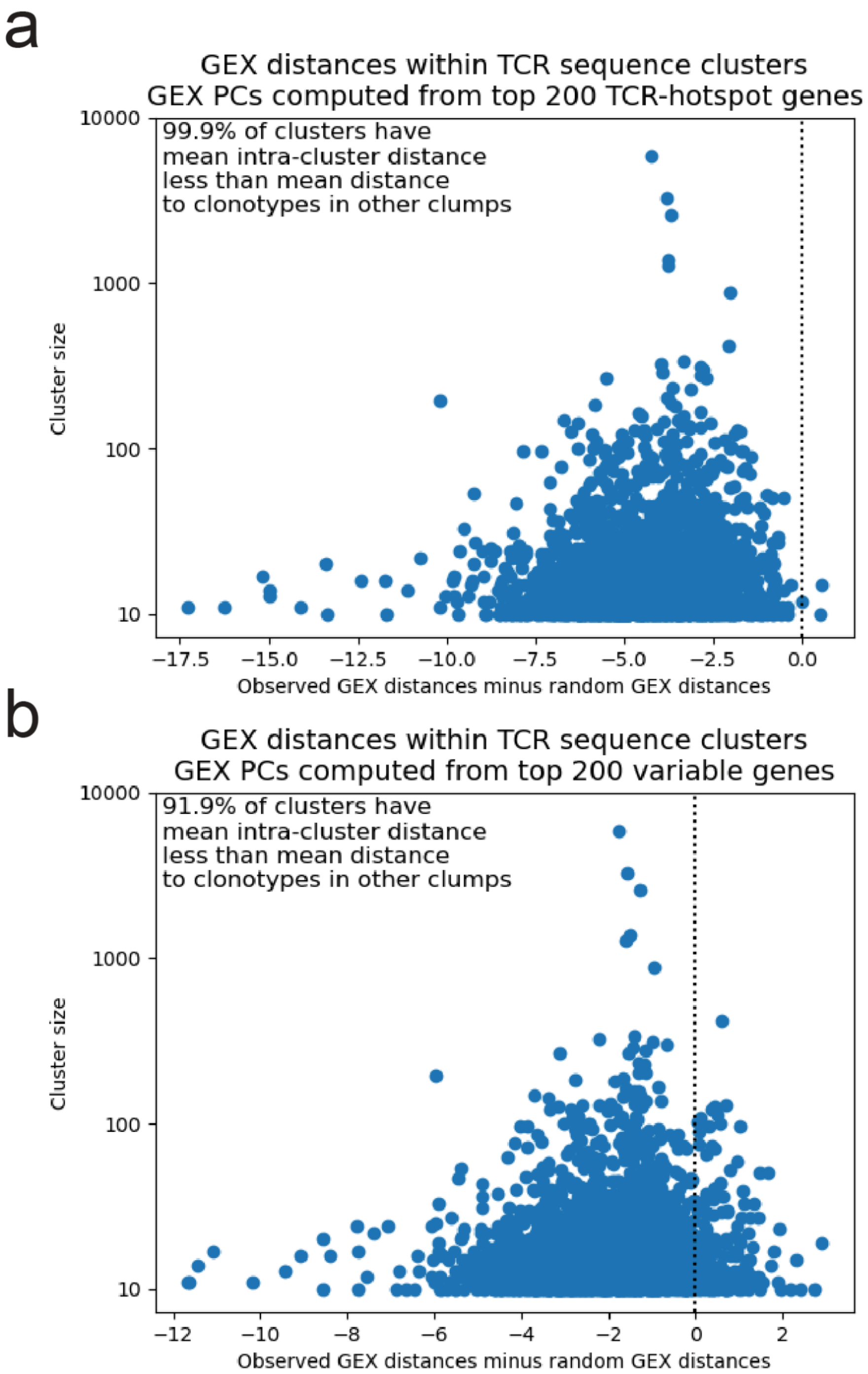


**Figure S4. Convergent TCR clusters have coherent gene expression profiles.** Clonotypes within TCR clusters are more similar to one another than to clonotypes in other clusters. Each dot represents a single TCR cluster (N=2173). The x-axis records the difference between mean intra- and inter-cluster clonotype GEX distances. (a) Clonotype GEX distance is calculated using the top 200 TCR-graph autocorrelated genes (TAGs; **Fig. S1b**). (b) Clonotype GEX distance is calculated using the top 200 highly-variable genes (HVGs).

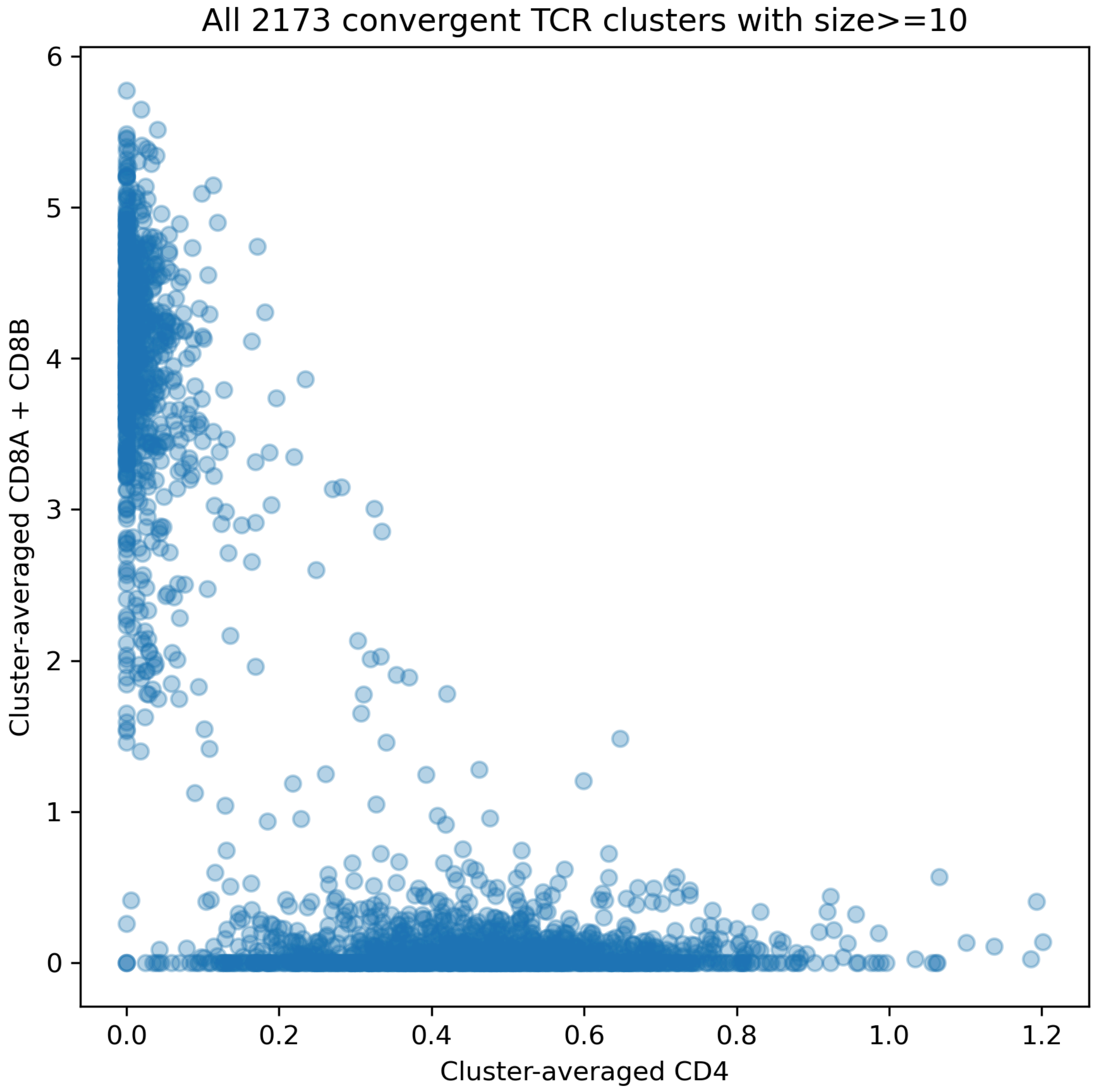


**Figure S5. Consistency of CD4 versus CD8 gene expression for convergent TCR clusters.** Each dot corresponds to one of the 2173 convergent TCR clusters of size >= 10, arranged by cluster-averaged CD4 expression (x-axis) and cluster-averaged CD8 expression (y-axis). The majority of the clusters show dominant expression of either CD4 or CD8.

### Pipeline for CDR3 AA bias analysis (main Figs. 3-5)

#### Step 1
##### Identify GEX neighbors

- Subset to CD4 (or CD8) cells
- Scale per-cell transcript counts to sum to 10000; log+1 transform; average over cells within expanded clonotypes
- Subset to the top 200 TCR-graph associated genes
- Run PCA, take top 20 PCs, compute Euclidean distances between clonotypes in PC space
- Identify K=5000 nearest neighbors for each clonotype

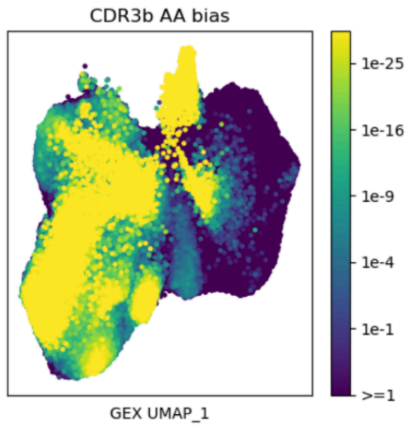


#### Step 2
##### Compute CDR3 amino acid frequencies in neighborhoods

- For each GEX neighborhood (defined by a central clonotype and its 5000 nearest neighbors), calculate the frequencies of each amino acid in the central regions of the CDR3a and CDR3b loops (separately) of the neighborhood TCRs
- Average these frequencies to derive a background profile

#### Step 3
##### Identify neighborhoods with biased AA frequencies

- Compare observed frequencies to expected frequencies (from the overall background profile) using chi-squared distribution
- Multiply the raw chi-squared $P$ value by the number of neighborhoods to correct for multiple testing
- Define significantly biased neighborhoods as those with adjusted $P$ value < $10^{-6}$

#### Step 4
##### Build landscape of biased neighborhoods

- Represent each biased neighborhood by its CDR3alpha and CDR3beta AA frequency profiles
- Run UMAP dimensionality reduction and Leiden clustering on these 40-dimensional vectors
- For each Leiden cluster, visualize the cluster-averaged AA frequency profile with a differential sequence logo

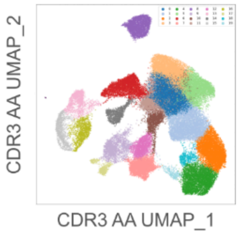


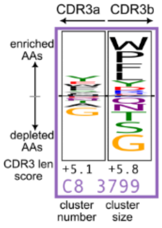


#### Step 5
##### Visualize TCR and GEX features mapped onto the landscape

- Standardize each TCR or GEX feature to have mean 0 and standard deviation 1 over all clonotypes
- Average the TCR or GEX features over each neighborhood and plot these neighborhood-averaged values on the 2D UMAP landscape

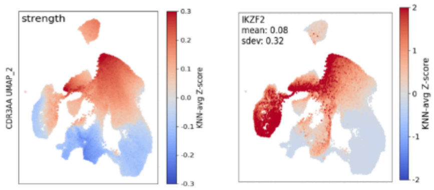


#### Step 6
##### Analyze gene expression in biased neighborhoods

- Use additional gene features to build a hi-res GEX landscape and clusters for the clonotypes with biased CDR3aa profiles
- Use gene expression, cohort metadata, and clonotype occurrence patterns for annotation

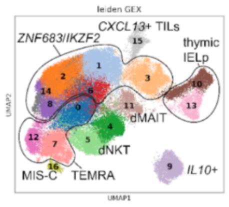


#### Step 7
##### Match neighborhood TCRs to literature TCR databases

- Match TCRs in biased neighborhoods to antigen-specific TCRs from published datasets w/ TCRdist
- Assess significance of matches using comparisons to a VDJ-shuffled background

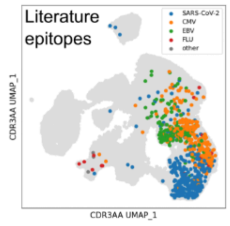

**Figure S6. Pipeline for CDR3AA bias analysis.** Details can be found in the Methods section.

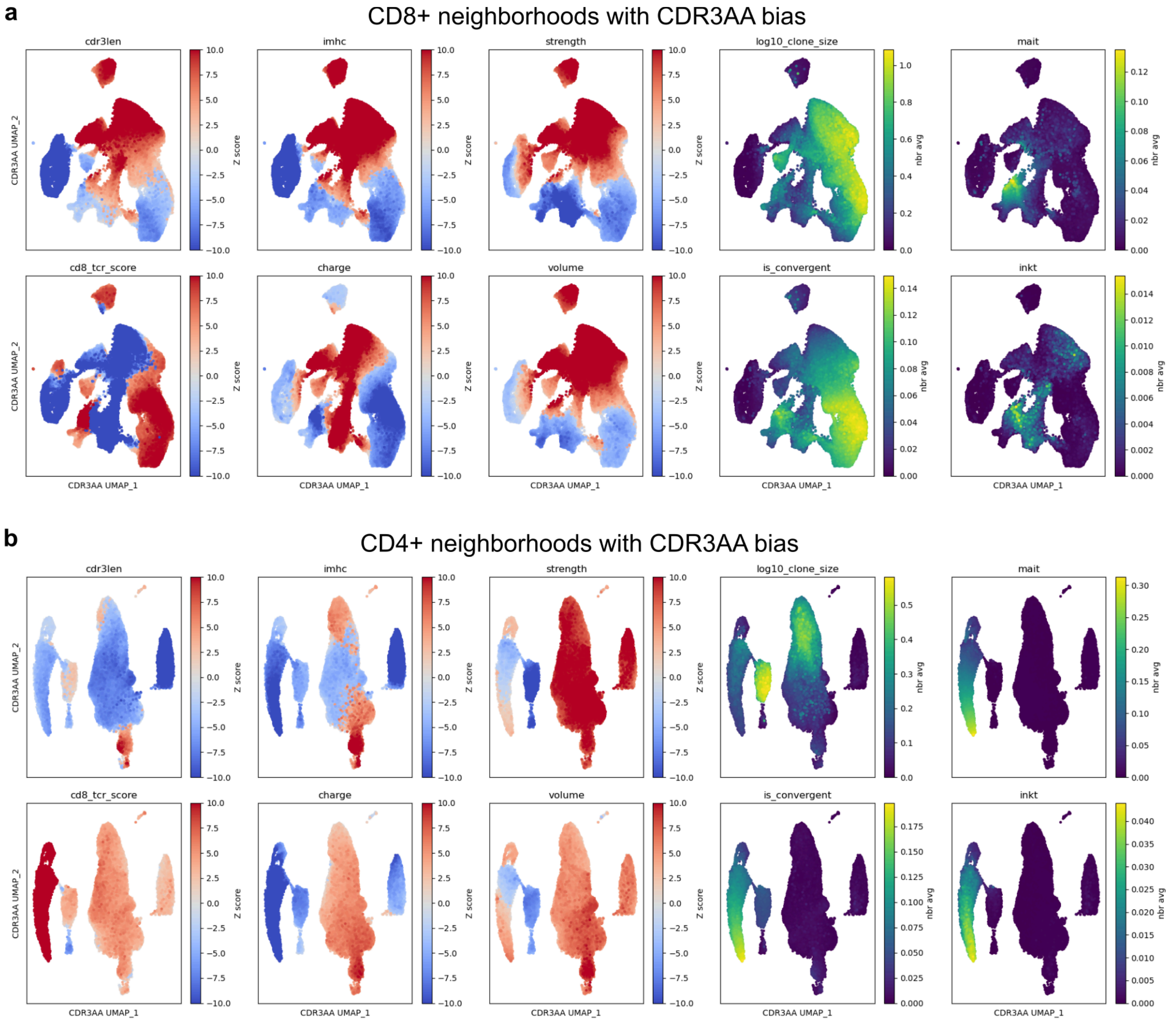


**Figure S7. CDR3AA bias landscapes colored by additional TCR sequence features.** Each dot corresponds to a CD8+ (panel a, N=71,528) or CD4+ (panel b, N=53,255) GEX neighborhood with significantly biased CDR3 amino acid composition. The dot is colored by the KNN-averaged value for the TCR feature indicated in the panel title. TCR features are defined in Ref (Schattgen et al. 2021). Briefly, 'cdr3len' measures the combined length of CDR3a and CDR3b; 'imhc' corresponds to the CoNGA iMHC score which captures sequence features (positive charge, increased volume and hydrophobicity, enrichment for cysteine) of a set of putatively MHC-independent TCRs; 'strength' assesses the frequency of the amino acids CFILMVW in the central regions of CDR3a and CDR3b; 'log10_clone_size' measures the degree of clonal expansion; 'mait' reflects matches to a MAIT sequence consensus defined by TRAV and TRAJ genes and CDR3a length; 'cd8_tcr_score' is a sequence based score optimized to discriminate CD8 from CD4 TCR sequences; 'charge' measures CDR3 amino acid charge in the central region (excluding the first and last four residues); 'volume' measures CDR3 amino acid volume in the central region; 'is_convergent' records whether a TCR belongs to one of the 2173 convergent TCR clusters of size 10 or greater; 'inkt' reflects matches to an iNKT sequence consensus defined by TRAV, TRAJ, and TRBV genes and CDR3a length. The scores in the left 6 panels were normalized by subtracting the mean and dividing by the standard deviation before neighbor averaging.

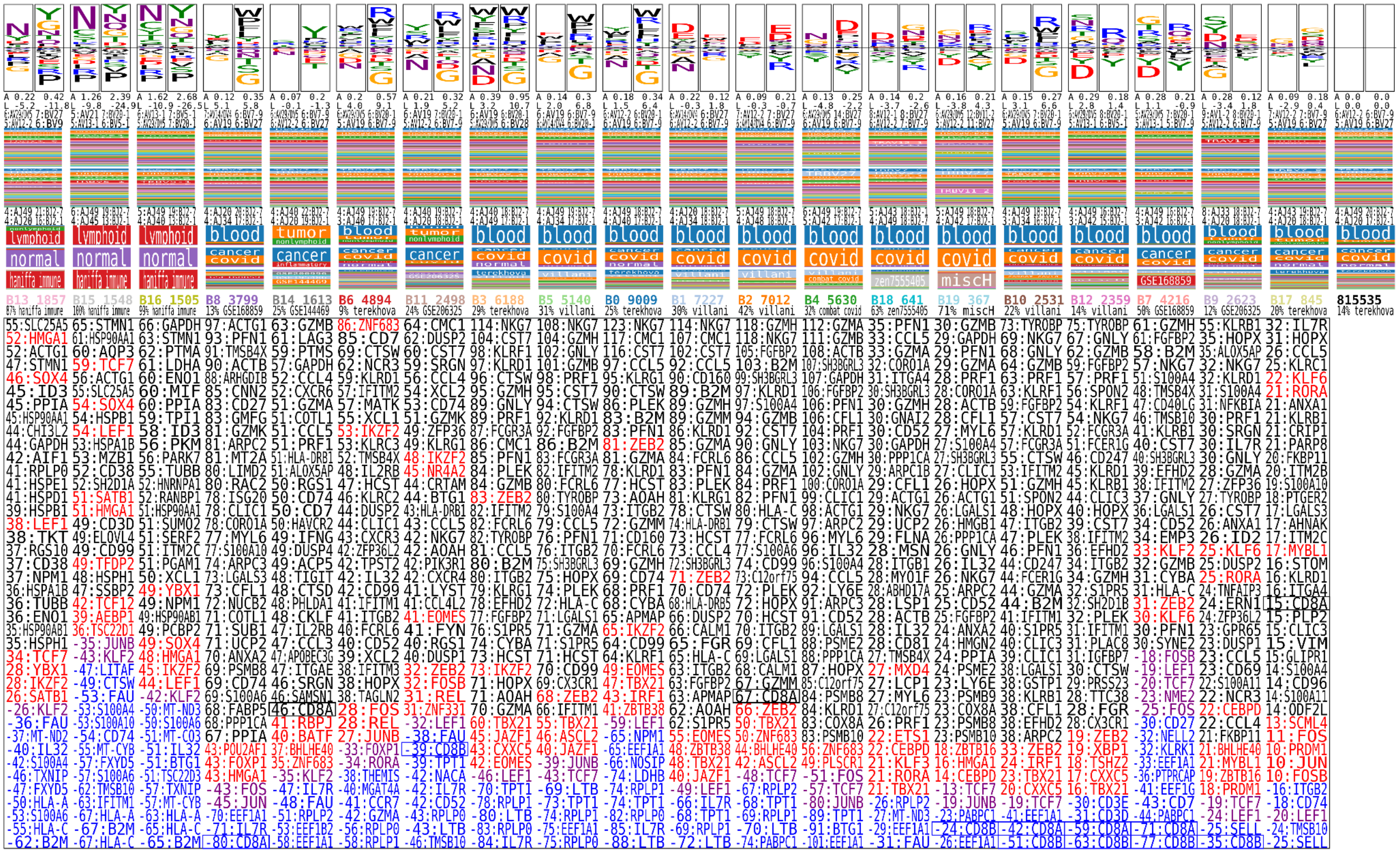

**Figure S8. Top differentially expressed genes for the CD8+ CDR3AA bias clusters.** Upregulated genes shown in black (red for transcription factors); downregulated genes shown in blue (purple for transcription factors).

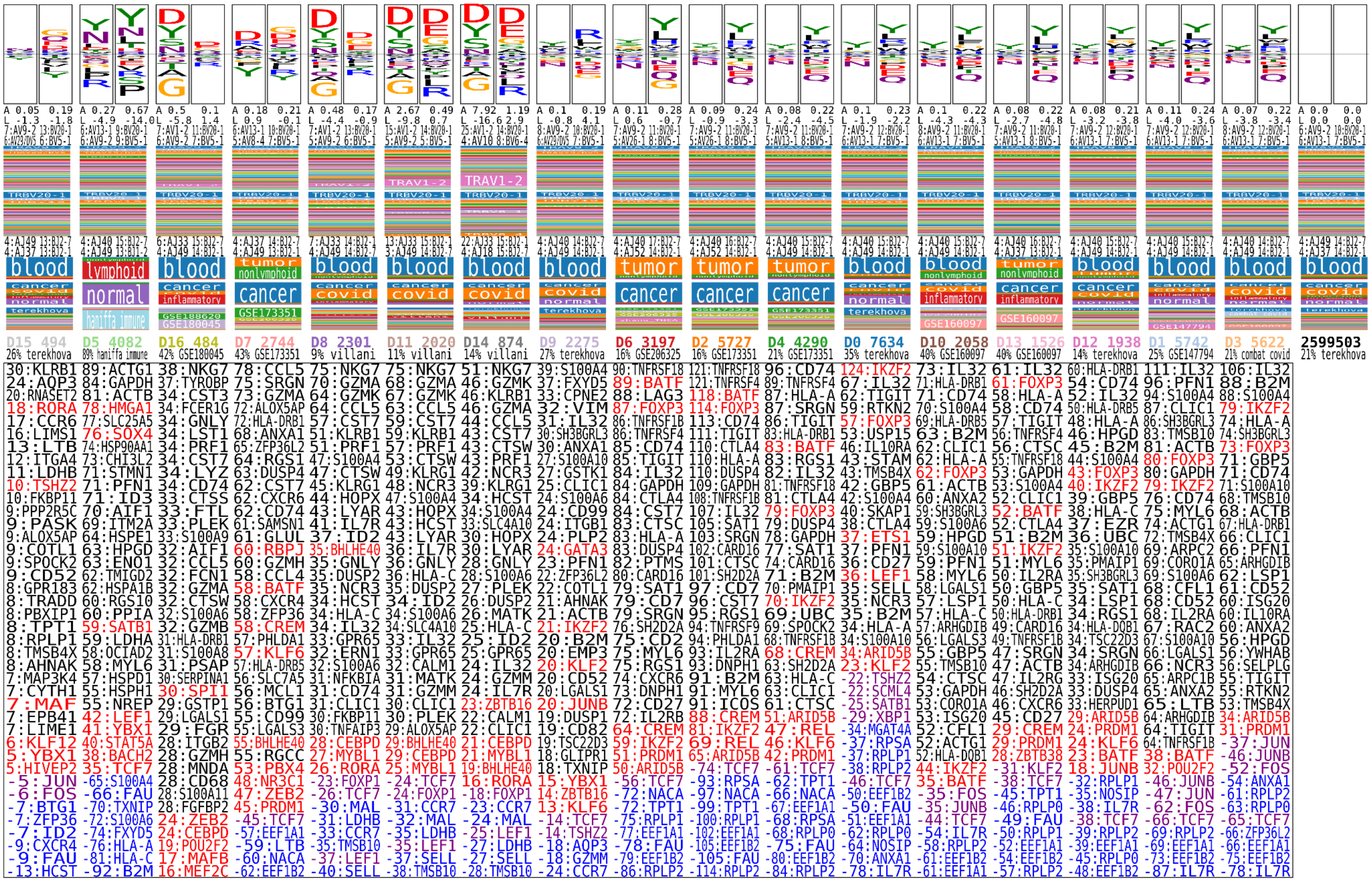

**Figure S9. Top differentially expressed genes for the CD4+ CDR3AA bias clusters.** Upregulated genes shown in black (red for transcription factors); downregulated genes shown in blue (purple for transcription factors).

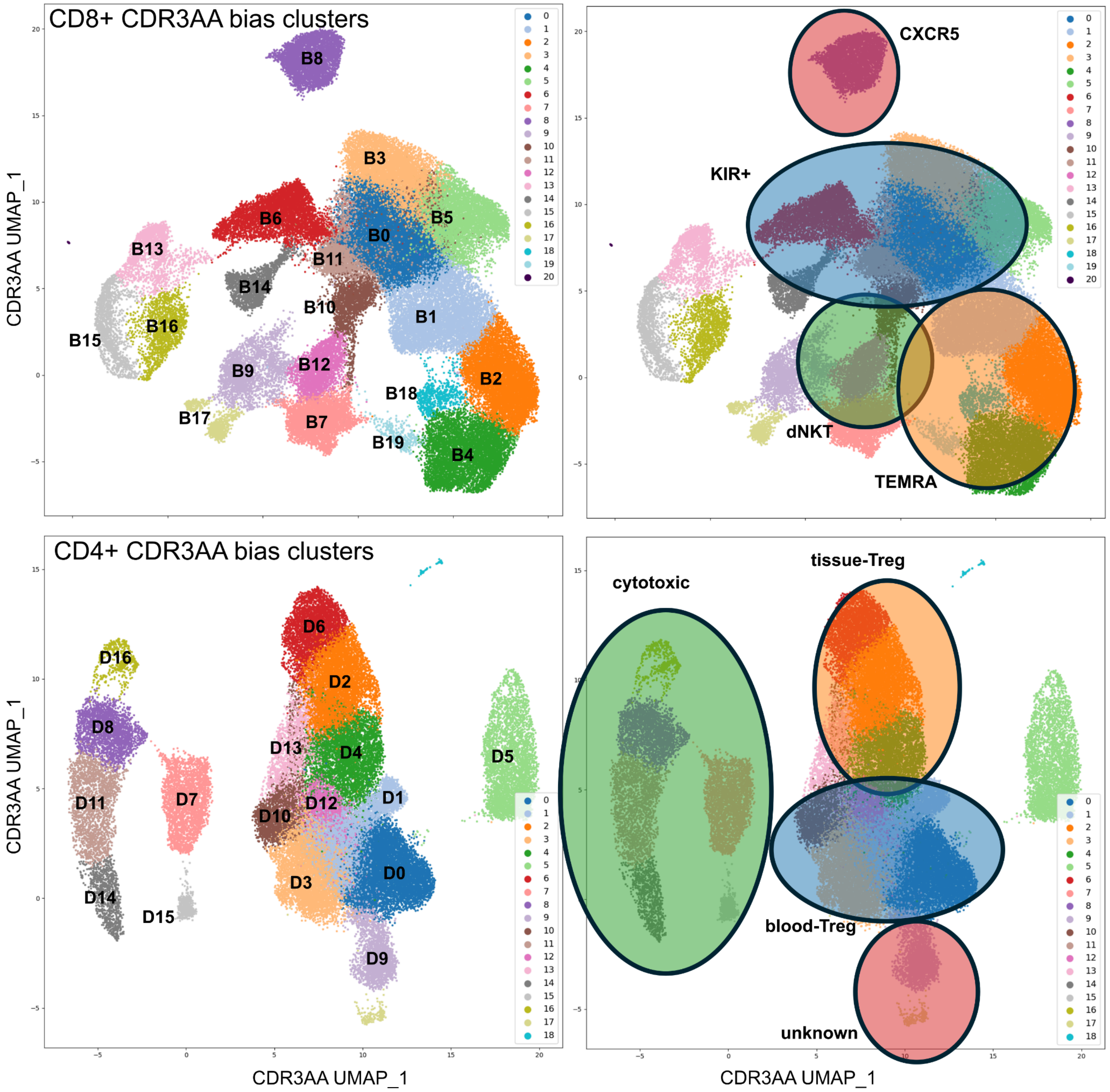


**Figure S10. Putative functional groupings of the CD8+ and CD4+ CDR3AA bias clusters as used in the bar plots in main text Figure 6.**

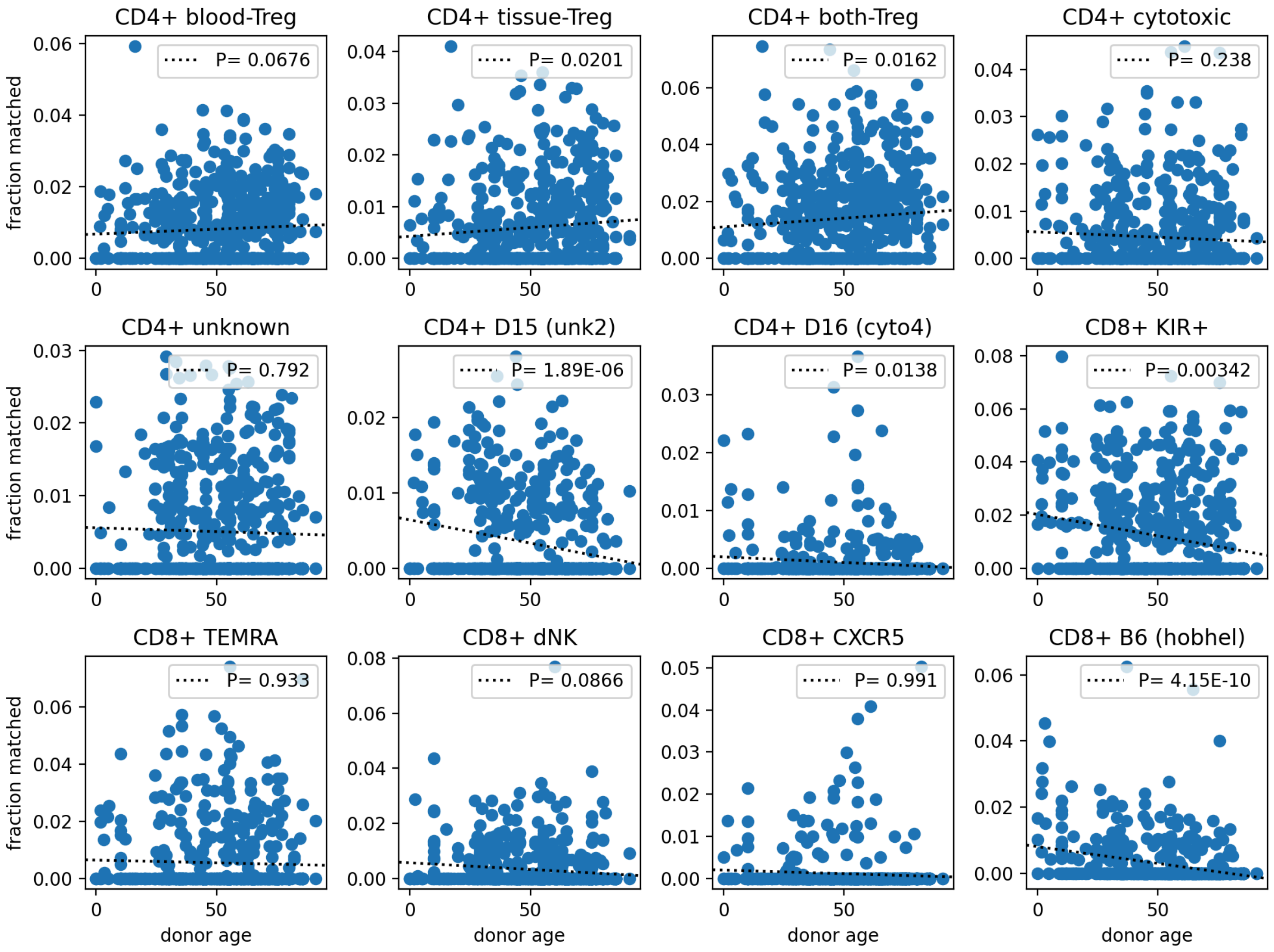


**Figure S11. CDR3AA-bias cluster matches as a function of donor age in the metaCoNGA dataset.** For each individual donor in the metaCoNGA atlas, clonotypes matching the CDR3AA bias clusters were identified as described in the main text ("Annotating new datasets with metaCoNGA features") by looking for overlap between high-scoring regions for the cluster-specific TCR and GEX scores. Panels show donor age (x-axis) plotted against the fraction of total clonotypes (y-axis) matching the cluster or cluster group (see **Fig. S10**) indicated in the panel heading. Correlations between age and matched fraction of clonotypes were calculated using a linear regression model that accounted for the total number of clonotypes (which declines with age and can also influence matching fraction). Correlation $p$-values are given in the panel legends. The three individual clusters (D15, D16, and B6) with the most significant correlations are plotted in addition to the 9 cluster groups.
